# Native extracellular matrix promotes human neuromuscular organoid morphogenesis and function

**DOI:** 10.1101/2023.05.19.541464

**Authors:** Beatrice Auletta, Lucia Rossi, Francesca Cecchinato, Gilda Barbato, Agnese Lauroja, Pietro Chiolerio, Giada Cecconi, Edoardo Maghin, Maria Easler, Paolo Raffa, Silvia Angiolillo, Wei Qin, Sonia Calabrò, Chiara Villa, Onelia Gagliano, Cecilia Laterza, Davide Cacchiarelli, Matilde Cescon, Monica Giomo, Yvan Torrente, Camilla Luni, Martina Piccoli, Nicola Elvassore, Anna Urciuolo

## Abstract

Human neuromuscular organoids (NMOs) derived from induced pluripotent stem cells (hiPSCs) hold a great potential to study (dys)functional human skeletal muscle (SkM) in vitro. The three-dimensional (3D) self-assembly of NMOs leads to the generation of spheroids, whose 3D organization cannot be controlled. Indeed, proper development, maturation and function of the innervated SkM require a well-defined multiscale 3D organization of the cells in a tissue-specific extracellular matrix (ECM) context. We hypothesized that extracellular structural imprinting along with hiPSC small-molecule-based differentiation could provide self-assembly guidance driving NMO morphogenesis and promoting the maturation and function of the human neuronal-coupled SkM in vitro models. We found that SkM ECM, provided as decellularized skeletal muscle, is able to reproducibly guide the morphogenesis of differentiating hiPSC toward multiscale structured tissue-like NMOs (t-NMOs). T-NMOs show contractile activity and possess functional neuromuscular junctions (NMJs), with mature neuromuscular system upon 30 days of hiPSC differentiation. We found that t-NMO could mimic altered muscle contraction upon administration of neurotoxins that act at NMJ level. Finally, we used hiPSCs derived from patients affected by Duchenne Muscular Dystrophy (DMD) to produce DMD t-NMOs that, upon neuronal stimulation, were able to mimic the altered SkM contractility and calcium dynamics typical of the disease. Altogether, our data confirm the ability of t-NMO platform to model in vitro human neuromuscular system (patho)physiology.

## Introduction

Organoids are 3D cell culture systems that have the ability to self-organize into aggregates forming tissue- or organ-like cytostructures^1^. In particular, human NMOs have been recently derived from both embryonic stem cells or hiPSCs, enabling the production of in vitro models containing both muscular and neuronal compartments to study functional human SkM^2–5^. The multicellular interactions between the neural and muscular systems represent one of the major advantages of such 3D in vitro models. Indeed, SkM homeostasis and function require a coordinate interaction between myofibers and motor neurons (MNs) at the NMJ, a complex chemical synapse formed between MNs and myofibers, responsible for central nervous system-controlled muscle contraction^3,6,7^. Studies based on the use of NMOs were able to recapitulate key aspects of myasthenia gravis^4^ and amyotrophic lateral sclerosis^2^. SkM organoids were also recently derived from hiPSCs to model myogenesis and muscle regeneration^5^. One major limiting step of organoids resides in the inability to control their self-assembly, which is in turn affecting reproducibility regarding cell type composition, architecture and functions^1^. In particular, the growth of NMOs into spheroids remains a major challenge for neuronal-coupled SkM modelling. In NMOs, the neuromuscular compartments are packed together within the spheroid and their self-assembly cannot be controlled. Therefore, how the functionality of the neuromuscular system model is affected by the uncontrollable spheric architecture cannot be predicted in currently available NMOs.

The innervated SkM is characterized by a multiscale 3D organization that guarantees its function^8–10^. This organization range from single to parallel bundles of anisotropic myofibers, associated to adult muscle stem cells and reached by single neural axons that sprout from nerves, toward coherent organization of muscular and neural cells within specific ECM domains^8–11^. It is well known that the proper development, maturation and function of the neuronal-coupled SkM also requires defined 3D organization of the cells in a context of structural and functional supporting ECM^3,8–11^. Indeed, a large body of studies demonstrated that ECM deposition structurally and functionally supports the 3D organization of myofibers in elongated and parallel bundles, which strongly impinge on their contraction ability and maturation^3,8–10^. Similarly, specific ECM components guide and sustain MN axons on the long way that separate their soma from their muscular targets, as well as for establishing and sustain NMJs, which in turn influence MN homeostasis and SkM function^11–13^.

Based on the active role of ECM in the neuromuscular system, we hypothesized that SkM-derived ECM could possess environmental imprinting cues that are able to guide hiPSC differentiation and morphogenesis toward functional NMOs that more closely resemble the in vivo multi-scaled organization of the tissue. To investigate this, we used native ECM, i.e. decellularized murine SkM (dSkM), as scaffolding biomaterial during hiPSC small molecule-mediated differentiation toward neuromuscular system. DSkMs are natural scaffolds that retain the native ECM mechanical integrity, biological activity, and 3D architecture of the tissue^14^. We previously showed that dSkMs support muscle regeneration in volumetric muscle loss models, including nervous system and NMJ formation^15–18^, and can be used in vitro to study myogenesis^17,19^, neural axon sprouting and to support the generation of neuromuscular 3D in vitro models^20^.

Here we show that direct differentiation of hiPSC in a context of SkM ECM by using dSkM allows the generation of reproducible multiscale structured t-NMOs, which more closely resemble the 3D organization of the neuronal-coupled SkM. This 3D morphological architecture was not achievable neither with hiPSC differentiated in standard 2D cell culture platforms nor in 3D self-assembled NMO condition. Interestingly, RNA-sequencing data revealed that the differential spatial organization of the hiPSC-derived neuromuscular system in vitro models correlates with differential gene expression profiles. By providing dSkM, we observed an improved maturity of the human 3D in vitro neuronal-coupled SkM model. Differentiated myofibers and SkM stem cells, as well as neural progenitors and differentiated MNs together with glial and Schwann cells, were identified in t-NMOs. Spontaneous contraction of t-NMOs was revealed starting from day 15 of hiPSC differentiation. High degree of SkM maturation was reached after 30 days of cell culture, with functional NMJs and fetal to adult acetylcholine receptor (AChR) subunit^21^ switch. Chemical or genetic approaches known to affect SkM contraction were implemented to confirm that t-NMOs could be used to reveal aberrant muscular functionality upon neural network stimulation. Neurotoxins that act at the NMJ were able to reduce or abolish muscle contraction upon neural stimulation of t-NMO. Moreover, t-NMOs derived from two different DMD patients displayed reduced contraction and altered calcium flux. Our results indicate that dSkM can be used to guide NMOs toward a multiscale 3D structure that mimic innervated SkMs in vitro, where the human neuromuscular system (patho)physiology can be studied.

## Results

### Decellularized SkM guide NMO morphogenesis

To reach our aim, dSkMs were derived from murine diaphragms as previously described^22^. We used diaphragm for its unique combination of myofiber innervation spatial distribution, together with reduced thickness for easy microscopic investigation^15,16^. Apart from the preservation of structural and topographical feature of native tissue^22^, we also confirmed that dSkMs preserved mechanical properties and partially retained the architecture of the neuronal network and AChR clustering of the native SkM tissue (Supplementary Fig. 1A-C). For hiPSC differentiation, we adapted a widely used protocol that is based on small molecules administration to recapitulate the embryonic developmental events that lead to myogenic commitment and later to myogenic terminal differentiation^23^. Notably, apart from SkM differentiation, such protocol has been shown to promote the generation of other non-muscular cell types in conventional 2D cell culture systems, including neurons and Schwann cells^24^.

To produce t-NMO, hiPSCs were seeded onto dSkM as single cells and differentiated for 30 days (Fig. 1A). Extensive 3D cell aggregates formed onto the scaffold from day 5 to day 10 of differentiation, and by day 30 of culture, cells completely invaded the available surface of dSkMs (Fig. 1B). To monitor initial hiPSC commitment, we performed whole mount immunofluorescence. Two days after cell seeding (day 0 of differentiation), hiPSCs were growing onto the dSkM as flat, compact epithelial colonies expressing octamer-binding transcription factor 4 (OCT4) and homeobox protein NANOG (Supplementary Fig. 2A). Two days after the induction of cell differentiation, the expression of the mesoderm-induction T-box transcription factor brachyury (TBRA), SRY-box transcription factor 2 (SOX2) and caudal type homeobox 2 (CDX2) was revealed, together with downregulation of NANOG, indicating the ability of dSkM to support hiPSC neuromesodermal progenitor commitment (Fig. 1C and Supplementary Fig. 2A,B). Reproducibility of such hiPSC-derived cell growth onto dSkMs was confirmed by multiple experiments (> 120 biological replicates, Supplementary Fig. 2C) and obtained with 5 different hiPSC lines along the entire study. In parallel with t-NMO production, the same differentiation protocol was applied to hiPSCs cultured onto standard 2D culture devices coated with commercial basement membrane-rich (Matrigel) or onto 3D Matrigel flat droplets to derive spheric NMOs (Supplementary Fig. 3). Of note, Matrigel droplets offer a 3D scaffolding natural biomaterial that does not contain structural, mechanical, topographical and compositional properties specific to SkM.

**Figure 1.**
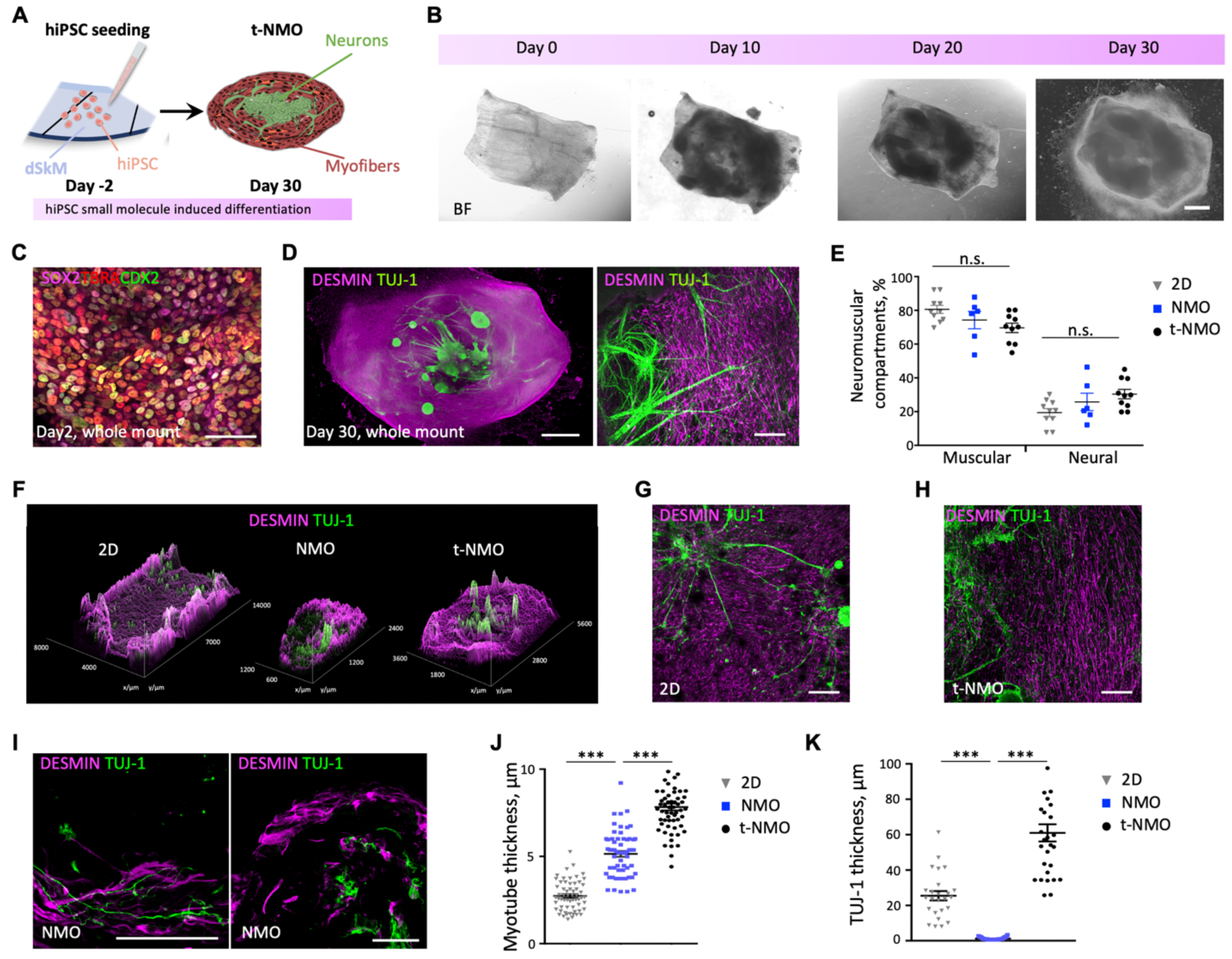
DSkMs support hiPSC-Neuromesodermal progenitor commitment and control t-NMO morphogenesis. **A.** Schematic illustration showing the strategy used for t-NMO generation. **B.** Representative bright field (BF) images showing morphological appearance of hiPSCs seeded onto decellularized muscles (Day 0) and undergoing neuromuscular differentiation after 10 (Day 10), 20 (Day 20) or 30 (Day 30) days. Scale bar, 2 mm. **C.** Confocal immunofluorescence imaging showing NMP committed cells (co-expressing SOX2, TBRA and CDX2) 2 days after hiPSC differentiation onto dSkM. Scale bar, 50 μm. **D.** Representative stereomicroscope (left) and confocal (right) immunofluorescence images of whole mount t-NMO after 30 days of differentiation stained for desmin (magenta) and TUJ-1 (green). Nuclei were counterstained with Hoechst (blue). Scale bars, 2 mm (left), 200 μm (right). **E.** Quantification of the area occupied by muscular (Desmin^+^) or neuronal (TUJ-1^+^) cells derived from hiPSCs after 30 days of differentiation into conventional cell culture plates (2D, gray), NMO (blue) or t-NMO (black). Data are shown as mean ± SEM of 6 to 10 independent replicates; one-way ANOVA with Tukey’s multiple comparisons test was used; n.s., not statistically significant. Statistical results are reported in Supplementary Table 1. **F.** Representative images showing desmin (magenta) and TUJ-1 (green) fluorescent signal spatial distribution in whole mount immunofluorescence stained 2D cultures and t-NMOs, and cross-sectioned stained NMOs. The coordinates indicate sample size in *xy* axis. **G-I.** Representative confocal Z-stack immunofluorescence images of 2D cultures (G), t-NMO (H) and NMO (I) stained for desmin (magenta) and TUJ-1 (green). Nuclei were stained with Hoechst (blue). Scale bars, 200 μm (G,H) and 50 μm (I). **J.** Quantification of desmin^+^ myotube cross-section (thickness) derived from hiPSCs after 30 days of differentiation in 2D cultures (gray), NMO (blue) or t-NMO (black). Data are shown as mean ± SEM of 3 independent replicates and dots represent single measured myotubes; one-way ANOVA with Tukey’s multiple comparisons test was used; \*\*\**P* < 0.001. Statistical results are reported in Supplementary Table 1. **K.** Quantification of TUJ-1^+^ neural projection cross-section (thickness) derived from hiPSCs after 30 days of differentiation in 2D cultures (gray), NMO (blue) or t-NMO (black). Data are shown as mean ± SEM of 3 independent replicates; dots represent single measured neural projections; one-way ANOVA with Tukey’s multiple comparisons test was used; \*\*\**P* < 0.001. Statistical results are reported in Supplementary Table 1.

According to the differentiation protocol applied^23^, and independently from the substrate used, 30 days after differentiation hiPSC-derived cells expressed muscle score genes (Supplementary Fig. 3B), as well as sets of genes belonging to neuronal cells (including neurons and Schwann cells) and mesenchymal cells (Supplementary Fig. 3C). Conversely, core genes of pluripotency were down-regulated (Supplementary Fig. 3D). Confirmation of the presence of neuronal and muscular cells was assessed by immunofluorescence for neuron-specific class III beta-tubulin (TUJ-1) and desmin^+^ myogenic cells (Fig. 1D and Supplementary Fig. 3E,F), respectively. Significantly higher percentage of myogenic area over the neuronal compartment was revealed in all 2D, NMO and t-NMO samples (Fig. 1E, Supplementary Table 1).

We used immunofluorescence analysis to evaluate compartmentalization and morphometric organization of the neuronal and muscular cells in the samples. Segregation of neuromuscular compartments was more evident in t-NMOs and NMOs, when compared to 2D samples (Fig. 1F). All the samples showed clusters of neural cell aggregates (TUJ-1^+^) that extended projections toward desmin^+^ myogenic cells (Fig. 1G-I and Supplementary Fig. 3E,F). However, we found that t-NMOs were characterized by myotubes and neural bundle projections with increased thickness, compared to both 2D cultures and NMOs (Fig. 1J,K). Longer and oriented myotubes, as well as longer neural projections were identified in t-NMO, when compared to 2D culture (Supplementary Fig. 3G-I).

This first set of data showed that dSkMs allowed direct hiPSC differentiation toward t-NMO, influencing NMO morphogenesis and leading to the generation of a bioengineered human NMO model that does not self-assemble in spheroid, more closely mimicking the native multiscale organization of the SkM.

### Morphological differences of the human neuromuscular in vitro models are paralleled by differential transcription profiles

To establish whether the morphological differences observed in t-NMO were paralleled by changes in the transcriptomic profiles when compared to 2D and NMO models, bulk RNA-sequencing analysis was performed in samples after 30 days of differentiation (Fig. 2, Supplementary Table 2 and 3). Three conditions were compared: cells differentiated in 2D, NMOs, and t-NMOs. This approach allowed us to investigate both the role of system three-dimensionality and the supportive action of dSkM on gene expression profiles.

**Figure 2.**
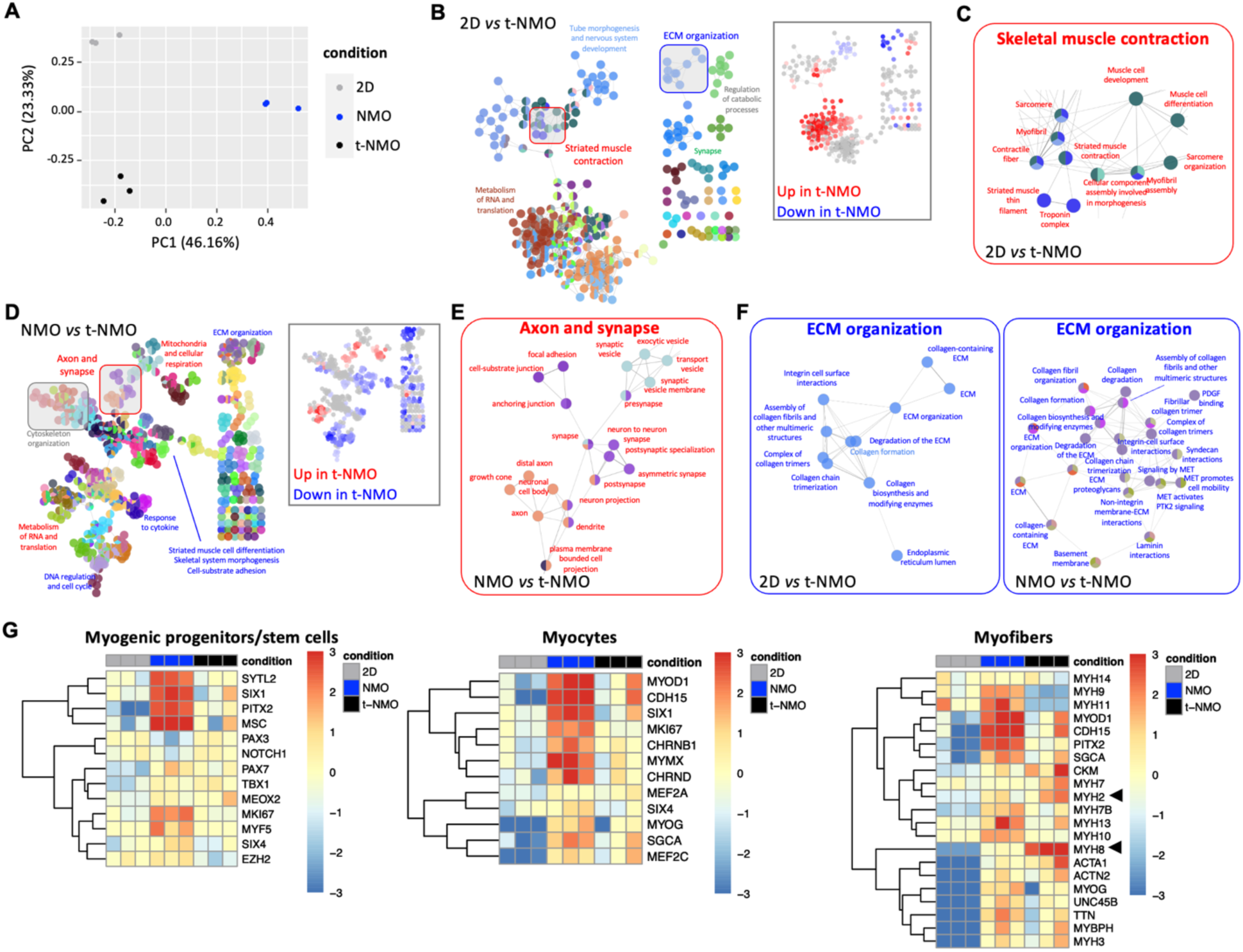
Neuromuscular system morphology organization is paralleled by differential gene expression and higher degree of maturation in t-NMO. **A.** Principal component analysis (PCA) of 2D, NMO and t-NMO samples. Each dot represents a single replicate. **B.** Enrichment analysis of differentially expressed genes between 2D samples and t-NMOs, within Gene Ontology and Reactome databases. Dot colors in the network highlight groups of categories with overlapping genes. Inset represents the same network where dots are colored according to log-fold change of gene expression. **C.** Enlargement of the portion of network highlighted in B, with categories related to SkM contraction, which are upregulated in t-NMO when compared to 2D samples. **D.** Enrichment analysis of differentially expressed genes between NMO and t-NMOs. Layout is similar to that in B. **E.** Enlargement of the portion of network highlighted in D, with categories related to neural axon and synapse formation, which are upregulated in t-NMO when compared to NMOs. **F.** Enlargement of the portions of network involved in ECM production highlighted in B (left) and D (right), whose categories are down regulated in t-NMO when compared to both 2D and NMO samples. **G.** Hierarchical clustering with heatmap visualization of the expression of manually annotated genes identifying myogenic stem cells (left), myocytes (middle) and myofibers (right). Arrows points at the adult MYH2 and neonatal MYH8 genes.

Principal component analysis (PCA) confirmed that 2D, NMO and t-NMO samples are transcriptionally well distinguishable (Fig. 2A), as confirmed by a high number of differentially expressed genes (DEGs), in the order of few thousands between pairs of conditions (Supplementary Table 2). With the aim to identify differentially expressed genes associated to the structural organization of the neuromuscular in vitro models, we first compared 2D samples with t-NMO, based on their similarity in neural axon sprouting and difference of myogenic organization. Interestingly, t-NMO showed strong upregulation of genes involved in SkM contraction and less clear differences in gene expression belonging to synapse-related categories, when compared to 2D neuromuscular models (Fig. 2B,C and Supplementary Table 3). We then compared NMOs with t-NMOs considering the muscular 3D organization of myofibers present in both the in vitro models and the differential neural projection organization. Accordingly, we found that t-NMO showed upregulation of genes related to neurotransmitters and synaptic function when compared to NMOs (Fig. 2D,E and Supplementary Table 3). As general feature, t-NMO showed downregulation of genes belonging to ECM production and remodeling, when compared to both 2D and NMO samples, suggesting an active contribution of the ECM provided by the dSkM in t-NMO (Fig. 2F). Moreover, t-NMO overexpressed genes associated with the categories of metabolism and RNA translation, when compared to both 2D and NMO samples (Fig. 2B,D and Supplementary Table 3). An increased level of maturity of myofibers in t-NMO was suggested by the analysis of the manually annotated gene list, which includes genes expressed by myogenic progenitors, myocytes and myofibers (Fig. 2G), when compared to both 2D and NMO samples. In particular, t-NMO highly expressed myofiber genes, including neonatal and adult myosin heavy chain (MYH) gene isoforms MYH8 and MYH2^25^, respectively (Fig. 2G).

Altogether, these data indicated that morphology and multicellular structural organization impinge on transcriptional profiles of the neuromuscular models, and that hiPSCs differentiated in the presence of native ECM increase the expression of genes involved in neuronal-coupled SkM maturation.

### Muscular and neural cell distribution in t-NMOs follows native-ECM organization

To validate the transcriptomic results, we focalized our subsequent studies on t-NMOs. Immunofluorescence analysis was used to confirm the presence of specific cell types belonging to muscular and neural compartments and reveal their spatial localization. Overall, t-NMOs showed a 3D organization of the multiple cell types, with neural clusters growing onto the dSkM, muscular cells present mainly in proximity of the dSkM reached by neural axons, and cells invading the dSkM scaffold (Fig. 1D, Fig. 3A and Supplementary Fig. 4A). Both paired box 7^+^ (PAX7^+^) stem cells and myogenin^+^ (MYOG^+^) differentiated cells were found in t-NMO (Fig. 3A,B and Supplementary Fig. 4B-C), confirming the preservation of both myogenic progenitors and differentiated cells. A significant higher ratio of PAX7^+^ cells was present in a non-proliferating state (Fig. 3C), and some PAX7^+^ cells located between pre-existing ghost myofibers and basal lamina, occupying the anatomical localization that defines satellite cells (Supplementary Fig. 4C). In agreement with transcriptomic data, the high degree of myofiber maturation was confirmed by the presence of myosin heavy chain (MYHC) expressing fibers reached by neural projections, and most of the nuclei identified in MYHC^+^ myofibers were expressing MYOG (Fig. 3D). We found that a consistent invasion of the ghost myofibers of dSkM by cells (Fig. 3E and Supplementary Fig. 4B-D), as well as myofibers located between the ghost fibers of the dSkM, showing typical striated pattern of myofiber cytoskeleton and expression of both slow and fast MYHC isoforms (Fig. 3D). Abundant basement membrane ECM component laminins were revealed in the proximity of the invaded dSkM (Fig. 3E). We confirmed that cell distribution within the dSkM followed the patterning imposed by the dSkM, as shown by quantification of the directionality of cells present in the proximity of ghost myofibers (Fig. 3F).

**Figure 3.**
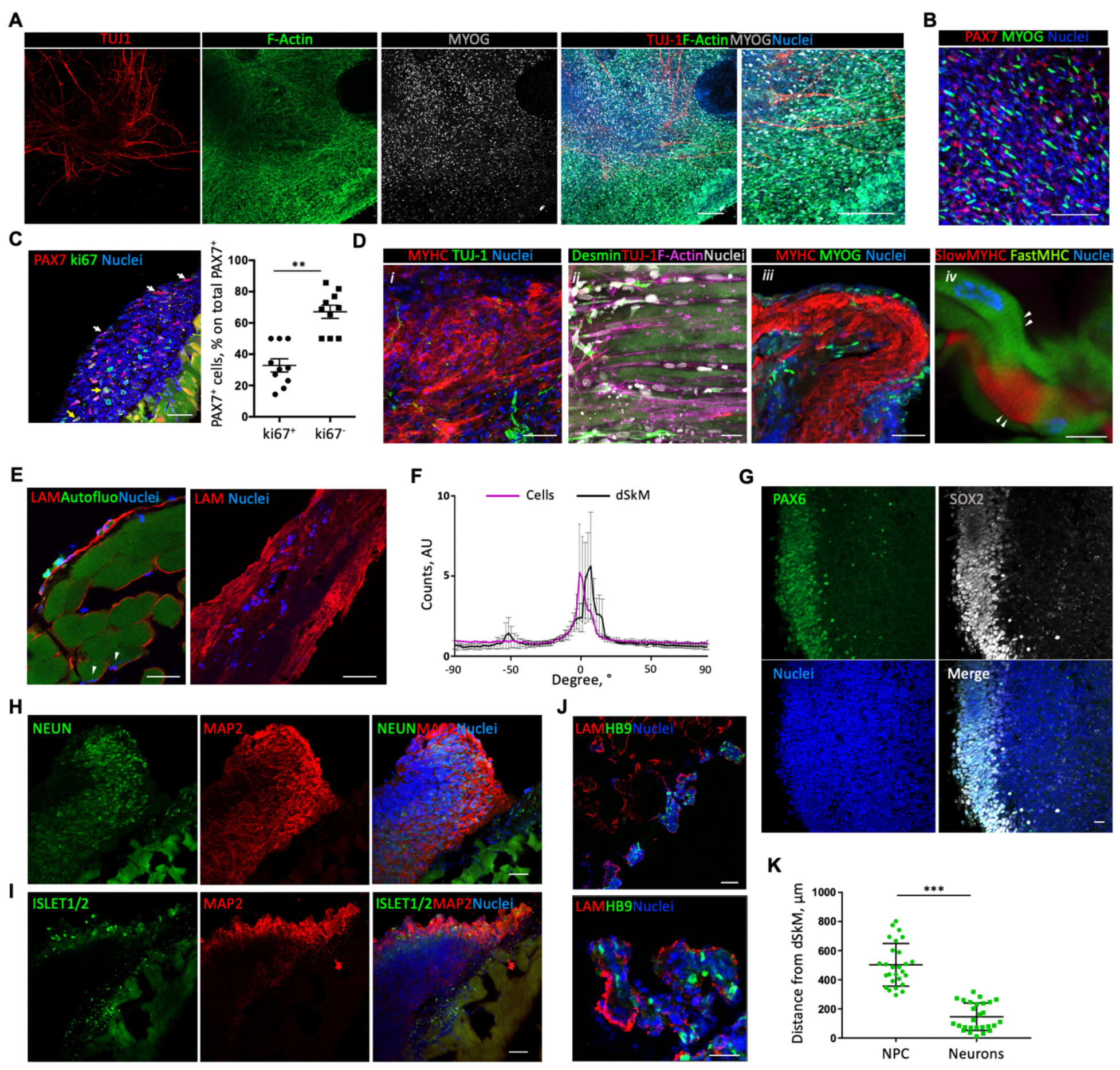
Native dSkM allows the generation of 3D t-NMOs with muscular and neural progenitors and terminal differentiated cells upon 30 days of differentiation. **A.** Representative confocal Z-stack images of the t-NMO stained for TUJ-1 (red), F-actin (green) and MYOG (grey). Scale bar, 200 μm. **B.** Representative images showing immunofluorescence staining for MYOG (green) and PAX7 (red) of t-NMO cryo-sections. Nuclei were stained with Hoechst (blue). Scale bar, 200 μm. **C.** Left panel, representative confocal Z-stack images showing immunofluorescence staining showing PAX7^+^ki67^-^ (white harrows) and PAX7^+^ki67^+^ (yellow harrows) cells. Scale bar, 50 μm. Right panel, quantification of proliferating (ki67^+^) and non-proliferating (ki67^-^) PAX7 positive (PAX7^+^) cells, expressed as percentage on the total PAX7^+^ cells. Data are shown as mean ± SEM of 3 independent replicates and dots represent single measured images; unequal variance Student’s *t*-test was used; \*\**P* = 0.0027. **D.** Immunofluorescence images showing myofiber maturation and dSkM invasion from cells. Representative confocal Z-stack images of t-NMO immunostained for *(i)* MYHC (red) and TUJ-1 (green); *(ii)* Desmin (green), TUJ-1^+^ neuronal extensions (red) and phalloidin (F-actin, magenta); *(iii)* MYHC (red) and MYOG (green); *(iv)* a higher magnification image showing myofibers expressing sarcomeric organization (white arrowheads) of fast and slow MYHCs. Nuclei were stained with Hoechst (blue or gray); Scale bars, 50 μm (*i, iii*), 25 μm (*ii*) and 10 μm (*iv*). **E.** Representative immunofluorescence confocal images showing deposited laminin (Lam, red) in proximity of nuclei (Hoechst, blue). Arrowheads point at nuclei located under laminin and in proximity of dSkM. Scale bars, 50 μm. **F.** Quantification of cellular and dSkM ghost myofiber directionality. Data are shown as mean ± SEM of 3 independent replicates. **G.** Representative confocal Z-stack images showing PAX6^+^ (green) SOX2^+^ (gray) neural progenitor cells localization. Scale bars, 20 μm. **H.** Representative confocal Z-stack images showing immunofluorescence staining of t-NMOs for the mature neuron makers NEUN (green) and MAP2 (red). Scale bars, 50 μm. **I.** Representative confocal Z-stack images showing immunofluorescence staining of t-NMOs for the MN markers ISL1/2 (green) and MAP2 (red). Scale bars, 10 μm. **J.** Representative confocal Z-stack images showing immunofluorescence staining of t-NMOs for the MN marker HB9 (green) and laminins (LAM, red). Scale bars, 50 μm (upper) and 20 μm (lower). **K.** Quantification of the distance of PAX6^+^SOX2^+^ neural progenitors (NPCs) or NEUN^+^ mature neurons from the dSkM in t-NMO cross-sections. Data are shown as mean ± SEM of 3 independent replicates and dots represent single measured cells; unequal variance Student’s *t*-test was used; \*\**P* < 0.0001.

We then investigated the neuronal compartment with the same approach. Neural progenitor cells (PAX6^+^ and SOX2^+^) were identified, together with mature neurons co-expressing neuronal nuclear protein (NEUN) and microtubule-associated protein 2 (MAP2; Fig. 3G,H). We also found clusters of neurons expressing LIM/homeodomain family of transcription factors ISLET1/2 (Fig. 3I), as well as surrounded by laminin and expressing the MN-determinant homeobox gene HB9, indicating the presence of mature MNs (Fig. 3J). Neural progenitor cells were identified in the upper layer of the neuronal clusters and in the more distant region with respect to the dSkM, when compared to mature neurons, that instead were identified on the borders of the clusters and in proximity of the contact with the dSkM (Fig. 3K). No cardiac nor smooth muscle cells were detected in the t-NMOs (Supplementary Fig. 4E,F), while we identified muscular and neuronal supporting cell types, as those expressing the myogenic fibroblast marker TE7 or the Schwann cells marker S100 calcium-binding protein B (S100β), respectively (Supplementary Fig. 3C and Supplementary Fig. 4G,H).

These data showed that t-NMOs contain multiple SkM and neuronal cell types with a complex 3D spatial organization that follows the structural architecture of the dSkM.

### T-NMOs contract and possess functional NMJs

In line with the above findings, we observed rhythmic and spontaneous contraction of t-NMO starting from day 15 of differentiation through the end of the experiment (Supplementary Video 1). Live imaging analysis of viable (calcein^+^) cells allowed to measure t-NMO contraction frequency (Fig. 4A), while calcium flux investigated via Fluo-4 imaging analysis identified physiological activity of the contracting area (Fig. 4B). Quantitative PCR gene expression analysis showed that the adult epsilon isoform^26^ was significantly upregulated in t-NMOs in respect to the fetal AChR gamma isoform, when compared to 2D cultures or spheric NMO (Fig. 4C and Supplementary Table 4). No spontaneous contraction was observed in 2D nor in NMO samples at comparable time points. Based on this observation, we then asked whether t-NMOs could possess functional NMJs. To investigate this, we first assessed the presence of NMJs by the investigation of pre- and post-synaptic elements via imaging analysis. By using fluorescent alpha-bungarotoxin (BTX) staining we could identify typical pretzel-like shaped pre-existing AChR clusters into the dSkM (Fig. 4D and Supplementary Fig. 1C), and smaller and digital-shaped newly formed AChR clusters located outside of the dSkM scaffold (Fig. 4D-F). We found that the ECM components known to support NMJ structural and functional integrity as collagen VI and perlecan were deposited in the proximity of AChR clusters (Supplementary Fig. 5B). Cells expressing S100β Schwann cell marker, particularly identified in the region of muscle-neural contact (Supplementary Fig. 4H), were also found in proximity of clustered AChRs contacted by neuronal extensions in t-NMOs (Supplementary Fig. 5C). The presence of nuclei and synaptic vesicle glycoprotein 2 (SV2) expressing neural structures located in the proximity of pre-existing AChR clusters of dSkM, suggested that the scaffold served as guidance for NMJ formation in the pre-existing patterned area (Fig. 4E). Moreover, NMJs were also found in areas distant from the dSkM, therefore identifiable of newly formed origin (Fig. 4F and Supplementary Fig. 5A).

**Figure 4.**
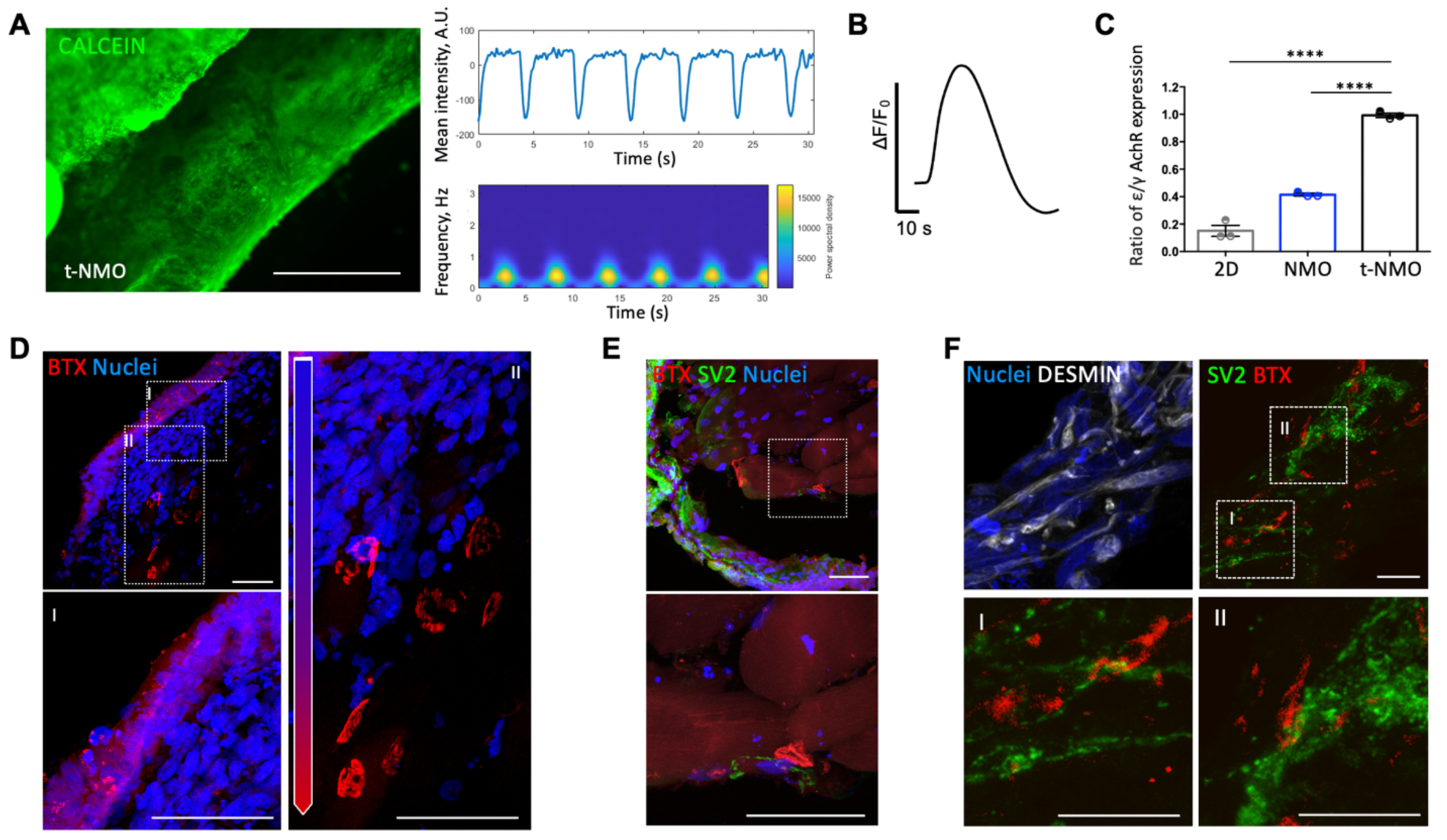
Native dSkMs support the formation of NMJs within a neural-coupled SkM model able to contract. **A.** Left panel, representative frame of live imaged (Calcein^+^) cells in t-NMO after 30 days of differentiation. Scale bar, 500 μm. Right upper panel, mean intensity variation registered during cyclic t-NMO contraction (A.U.). Right bottom panel, relative quantification of contraction frequency (Hz) registered during the time (s) represented by the heat map. **B.** Representative peak amplitude spectrum (ΔF/F_0_) of t-NMO basal activity detected with Fluo-4 live imaging analysis 30 days after differentiation. **C.** Ratio of gene expression between AChR adult (χ) and fetal (ψ) isoforms, comparing 2D (grey), NMO (blue) and t-NMO (black). Data are normalized for myosin heavy chain gene expression and are shown as mean ± SEM of 3 independent replicates. One-way ANOVA with Tukey’s multiple comparisons test; \*\*\*\**P* < 0.0001. Statistic is reported in Supplementary Table 4. **D.** Representative immunofluorescence images showing α-bungarotoxin^+^ (BTX, red) regions in t-NMO at day 30 of differentiation. Insets I and II show smaller and digital-shaped newly formed AChR clusters located out from the dSkM scaffold and pre-existing AChR clusters into the dSkM, respectively. Nuclei were stained with Hoechst (blue). Scale bar, 50 μm. **E.** Representative immunofluorescence images of BTX^+^ regions (red) in close contact with SV2 (green) in t-NMO at day 30. Nuclei were stained with Hoechst (blue). Scale bar, 50 μm. **F.** Representative immunofluorescence images of BTX^+^ AChR clusters (red) in desmin-positive myotube-enriched regions (white), found in close contact with SV2 neural projections (green) in t-NMO at day 30. Insets I and II show higher magnification of such contacts. Nuclei were stained with Hoechst (blue). Scale bar, 50 μm.

To confirm that the above NMJs were functional, live imaging analysis was performed to evaluate both contraction and calcium flux activity of the muscular compartment upon t-NMO stimulation with glutamate (Glu, Fig. 5A). We used acetylcholine (ACh) administration as controlled activity of myofiber responses, and whole mount immunofluorescence post-analysis was performed on the same biological replicates to identify the neural and muscular areas. We found that t-NMO contracted and showed calcium transients upon administration of both Glu and ACh (Supplementary Video 2 and Fig. 5B). Altogether, these data indicate that dSkM sustains the formation of functional and mature NMJs into neuronal-coupled muscle models that are able to contract.

**Figure 5.**
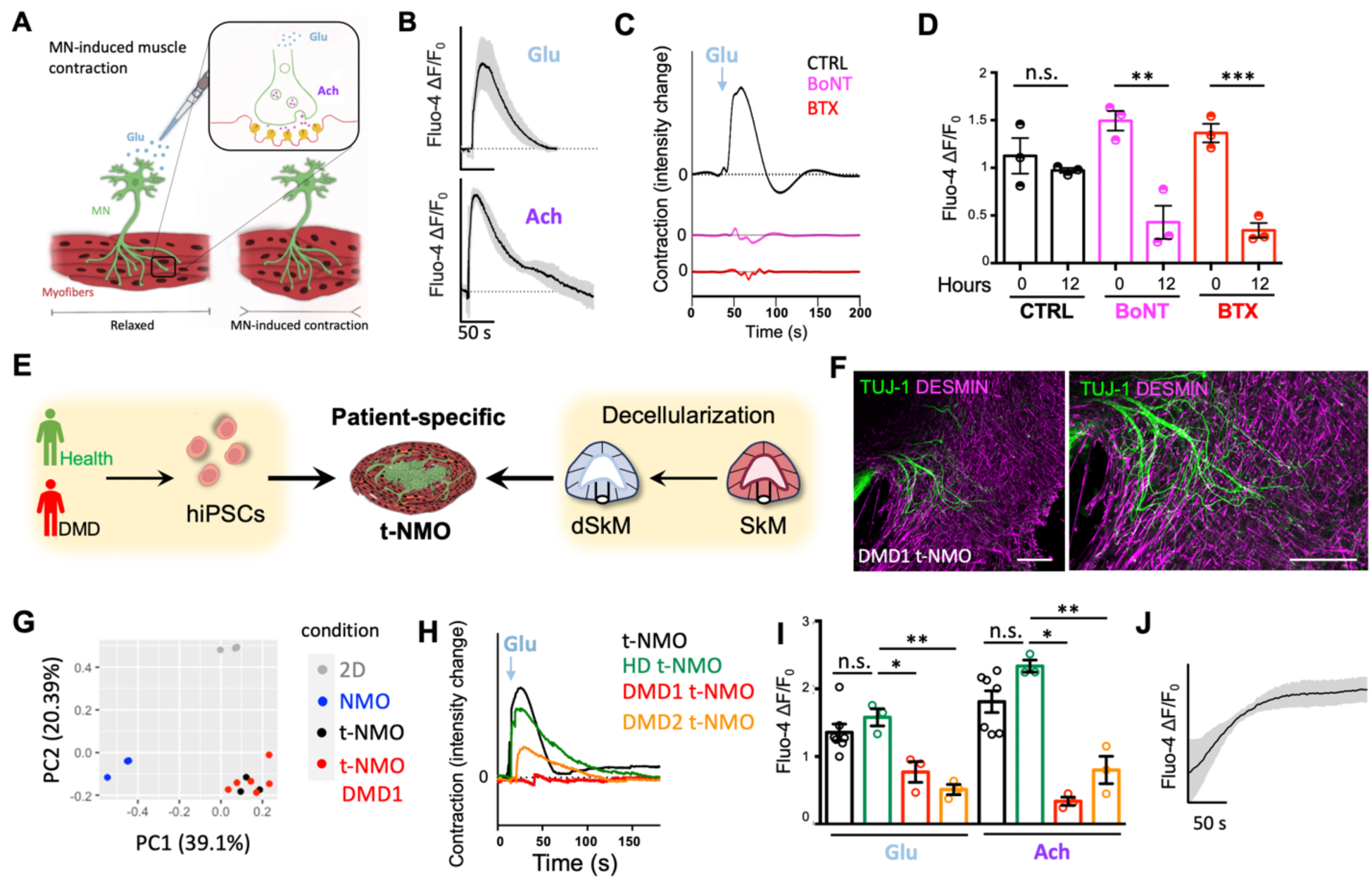
T-NMO reproduce (dys)functional activity of human SkM upon neuronal stimulation. **A.** Schematic illustration showing the strategy used for testing NMJ function in t-NMO. Glutamate (Glu) was supplemented to induce MN-mediated release of acetylcholine (ACh) and muscle response via NMJs. **B.** Representative average of peak amplitude spectrum (ΔF/F_0_) in t-NMOs upon Glu (upper panel, n = 3) or ACh (lower panel, n = 5) stimulations detected with Fluo-4 live imaging analysis 30 days after differentiation. ACh supplementation was used as control for muscle responses. Dotted lines correspond to the baseline equal to 0. **C.** Representative quantification of contraction of t-NMO differentiated for 30 days and stimulated with Glu after 12 hours of treatment with BoNT or BTX. As control, t-NMOs not treated with neurotoxins were analyzed (CTRL). Data are shown as mean ± SEM of 3 independent replicates. Dotted lines correspond to the baseline equal to 0. **D.** Quantification of calcium peak amplitude (ΔF/F_0_) detected with Fluo-4 live imaging analysis upon Glu stimulation of t-NMOs before (0 hours) or after 12 hours of treatment with BoNT or BTX. As control, t-NMOs not treated with neurotoxins were analyzed at 0 and 12 hours (CTRL). Data are shown as mean ± SEM of 3 independent replicates. One-way ANOVA with Tukey’s multiple comparisons test, n.s. not significant, \*\**P* = 0.0006, \*\*\**P* = 0.0009. Statistical results are reported in Supplementary Table 5. **E.** Schematic illustration showing the strategy used for producing patient specific t-NMO form healthy donors (Health) or Duchenne muscular dystrophy patients (DMD). **F.** Representative confocal Z-stack immunofluorescence images of DMD1 t-NMO stained for desmin (magenta) and TUJ-1 (green). Nuclei were stained with Hoechst (blue). Scale bars, 200 μm. **G.** Principal component analysis (PCA) of 2D (grey), NMO (blue) and t-NMO samples from BJ-hiPSCs (black) and DMD1 t-NMOs (red). Each dot represents a single replicate. **H.** Representative quantification of live imaging analysis showing contraction upon Glu stimulation of t-NMO (black), HD t-NMO (green), DMD1 t-NMO (red) and DMD2 t-NMO (orange) after 30 days of differentiation. Dotted lines correspond to the baseline equal to 0. **I.** Quantification of calcium peak amplitude (ΔF/F_0_) detected with Fluo-4 live imaging analysis upon Glu and ACh stimulation of t-NMO (black, n = 7), HD t-NMO (green, n = 3), DMD1 t-NMO (red, n = 3) and DMD2 t-NMO (orange, n = 3) after 30 days of differentiation. One-way ANOVA with Tukey’s multiple comparisons test, n.s. not significant; for Glu stimulation, \**P* = 0.001, \*\**P* = 0.0023; for Ach stimulation, \**P* < 0.0001, \*\**P* = 0.0007. Statistical results are reported in Supplementary Table 7. **J.** Representative average of peak amplitude spectrum (ΔF/F_0_) in DMD1 t-NMOs upon Glu stimulation detected with Fluo-4 live imaging analysis 30 days after differentiation (n = 3).

### Altered muscular responses can be observed upon MN stimulation in t-NMO that mimic pathologic conditions

After we thoroughly characterized the morphological and functional features of t-NMOs, we set up two experimental strategies to understand if t-NMOs could reveal altered muscular physiology upon MN activation and after only 30 days of hiPSC differentiation. At first, we tested whether MN-mediated muscle contraction was altered in presence of toxins known to block NMJ functionality. Specifically, we used botulinum neurotoxin A (BoNT) and BTX to induce pre-synaptic^26^ or post-synaptic^27^ blocks of the NMJ, respectively. Both toxins are expected to inhibit the muscle activity upon glutamate-mediated MN stimulation^26,27^. To do so, t-NMO contraction was measured via live imaging analysis before and after 12 hours of treatment with BoNT or BTX. As a control, we analyzed t-NMO contraction in absence of toxins at the same time points (0 and 12 hours). We found that only in presence of toxins, NMJ-mediated contraction was reduced when compared to untreated samples (Fig. 5C). Accordingly, a significant reduction of calcium flux was observed in the muscular areas upon Glu stimulation in both BoNT and BTX treated t-NMOs, when compared to untreated samples (Fig. 5D and Supplementary Table 5). By mimicking altered neuronal-induced muscular activity, these data demonstrated that t-NMO correctly responds to neurotoxin-driven NMJ block.

Finally, as proof of principle for t-NMO efficacy in modeling the pathophysiology of human neuronal-coupled SkM in vitro, we derived t-NMO from healthy donors and Duchenne muscular dystrophy (DMD) patients (Fig. 5E). DMD is a neuromuscular disease due to mutations in the dystrophin gene, characterized by a progressive muscle-wasting condition^21^. Dystrophin connects the contractile apparatus inside the myofiber to the ECM, and is enriched at the NMJ^29–31^. Here the aim was to understand if by provoking MN-induced muscle contraction, we could observe pathologic events in the muscular counterparts of DMD t-NMO that could mimic known DMD muscular phenotypes, as reduced muscle contraction and calcium dysregulation^28–31^.

We used the fast and highly efficient method of microfluidic cell reprogramming technology^32^ to initially generate DMD1 hiPSCs starting from muscular primary cells of a DMD patient carrying a stop codon mutation that lead to the loss of dystrophin production^33^ (Supplementary Fig. 6A). We then applied the same myogenic differentiation protocol to derive DMD1 t-NMOs. In accordance with previous results, DMD1 t-NMOs showed reproducible morphogenesis of tissue-like compartmentalized muscular and neuronal differentiated cells, that showed similar proportion compared to t-NMOs (Supplementary Fig. 6B-D). Bundles of myogenic cells were reached by elongated and structured neuronal projections (Fig. 5F). We confirmed absence of dystrophin protein in the muscular compartment and presence of NMJ upon 30 days of differentiation in DMD1 t-NMO (Supplementary Fig. 6E,F). Bulk RNA-seq analysis was performed to evaluate the overall gene expression profile of the DMD1 t-NMO. PCA analysis showed that DMD1 t-NMO clustered together with dystrophin expressing t-NMO (Fig. 5G), reinforcing the concept of high reproducibility of our model. Only 34 DEGs were up-regulated in t-NMO and 54 down-regulated compared to t-NMO DMD samples (Supplementary Table 6). Importantly, altered contraction activity was observed in DMD1 t-NMOs upon MN stimulation with glutamate, with significant lower degree of displacement, when compared to t-NMOs (Supplementary Fig. 6G). Based on these results, we then produced t-NMOs by using hiPSCs derived from another DMD patient (DMD2) and one healthy donor (HD) primary culture (Supplementary Fig. 7A,B). We therefore derived t-NMO from DMD2 and HD hiPSCs, confirming the expected morphogenesis and neuromuscular system organization upon 30 days of differentiation, and reproducing the proportion between neural and muscular compartments observed with the other hiPSC lines (Supplementary Fig. 7C-E). Importantly, we found that upon MN-induced glutamate stimulation both DMD1 and DMD2 t-NMOs showed reduced contraction, when compared to both HD t-NMOs and the control t-NMOs (derived from BJ-hiPSCs; Fig. 5H). Moreover, significantly reduced calcium spikes were observed in DMD1 and DMD2 t-NMOs, when compared to HD t-NMOs and the control t-NMOs upon both Glu and ACh stimulations (Fig. 5I and Supplementary Table 7), together with altered calcium handling upon MN-induced glutamate stimulation (Fig. 5J).

Altogether, these data reinforce the hypothesis that native ECM can reproducibly guide NMO morphogenesis, allowing the generation of multi-scaled organized t-NMO able to mimic SkM dysfunctionality.

## Discussion

There is an urgent request to increase our knowledge on human neuromuscular system (patho)physiology to finally improve the translation of preclinical studies to medicine. The recently reported NMO^2,4^ derived from hiPSC-derived neuromesodermal progenitors opened a new promise for studying the human neuromuscular system (dys)functionality. NMO technology is based on the generation of uncontrollable self-assembling spheric aggregates, that lack the multiscale 3D organization of myofibers and associated MN axons observed in vivo. This limit is of great relevance when the functionality of the neuromuscular system needs to be investigated in vitro. From the anisotropic 3D organization of SkM cells towards increased complexity of multicellular components of a functional SkM, 3D organization is a prerequisite for the function of innervated SkM^34^. The generation of a multiscale structured 3D NMO from hiPSCs has to our knowledge not been described yet. The use of bioengineering approaches to control organoid morphogenesis for consistent reproducibility and functionality represents a step forward in the translation of organoid technology^35,36^.

Here we developed a bioengineering strategy to derive multiscale structured NMOs, by combining the guidance cues given by the native SkM ECM with hiPSC technology. We found that by providing dSkM during hiPSC differentiation, NMO morphogenesis could be controlled. Using this strategy, we were able to produce human 3D innervated-SkM in vitro models (t-NMO) that better resemble the in vivo organization of the neuromuscular system without any pre-differentiation step of hiPSCs. As shown for NMO^2,4^, neural and muscular tissue compartmentalization was observed also in t-NMOs. However, the complexity of both muscular and neural axon bundle organization, as well as their interacting areas, were greater in t-NMO. Moreover, the specific neuromuscular morphology observed in t-NMOs was paralleled with differential transcription profiles, indicating greater level of neuromuscular maturity in guided t-NMO, when compared to self-assembled samples. According with other studies^34^, we confirmed that three-dimensionality improved myogenic compartment maturation. Indeed, improved maturation of myofibers was observed in both NMO and t-NMO, when compared to 2D cultures. On the contrary, neural maturation and synapsis formation were promoted in 2D cultures and t-NMOs, suggesting that the increased neural axon sprouting, likely related to the available 3D space for axonal growth of such culture conditions, could be involved in the maturation process. The supplied dSkMs supported hiPSC differentiation in terminal differentiated, progenitor and supporting stromal neural and muscular cells. Myogenic maturation in spheric NMOs was reached with longer culture time^2,4^. Both mechanical properties and ECM composition are known to influence stem cell differentiation and organoid morphogenesis^35–37^. Here we found that dSkM preserved mechanical properties of native SkM, and we previously showed that topography and native ECM composition of the native tissue were maintained upon decellularization^15–18,38^. Therefore, we can hypothesize that dSkM has an active role in hiPSC differentiation and/or in NMO morphogenesis, and finally in neuromuscular maturation. Two studies showed that decellularized tissues promoted hiPSC differentiation towards tissue-matched cell types, such as hepatocytes^39^ or endothelial cells^40^. Future studies aimed to dissect the molecular mechanisms involved in t-NMO formation could increase our knowledge on the role played by ECM in driving cell differentiation and morphogenesis in human tissue, as well as clarify the biology that governs the self-assembling of organoids. At increasing level of complexity, we also found that aside serving as a scaffolding biomaterial onto which NMO could grow, cells derived from hiPSCs also invaded pre-existing histological areas of dSkMs. Such *ghost* anatomical areas preserved in the dSkMs were invaded by anatomical matched-committed cells, such as those that repopulated dSkM myofibers or localized in satellite cell position or colonized pre-existing murine AChR clusters. These results are in agreement with previous in vitro and in vivo studies, where adult stem cells or adult progenitors were shown to invade and repopulate the *ghost* dSkM with anatomical matched-committed cells^15–18^. Altogether these evidences support the hypothesis that dSkM could also have a role in cell homing. Further studies in this direction can open up new strategies for engineering human organoids.

In line with our hypothesis, the morphological organization of t-NMO was also associated with increased functional activity of the SkM, as t-NMO shows spontaneous contraction starting from 15 days of differentiation. In t-NMOs, MNs and myofibers interact to generate functional and mature NMJs within a supporting ECM network in only 30 days of differentiation. Indeed, we were able to experimentally trigger SkM contraction by administering MN or myofibers specific neurotransmitters to the culture media, measuring SkM physiological activity in terms of contraction and calcium dynamics.

By containing functional components of the human neuromuscular system, together with high reproducibility and microscopy-friendly investigation for functional readouts, t-NMOs represent an interesting tool for studying neuromuscular dysfunctionality. As proof of principle, we set-up two experimental strategies to model SkM dysfunctionality. With the first approach, we used neurotoxins known to act at the NMJ to measure altered SkM physiology upon MN stimulation. As reported in literature^26,27^, treatments of t-NMOs with BoNT and BTX neurotoxins were inhibiting SkM contraction resulting from Glu-mediated MN stimulation. This confirmed the ability of t-NMO to be used to study in vitro neuronal-mediated SkM (patho)physiology. Finally, we modeled DMD, a disease caused by mutations in the X-linked *DMD* gene associated with SkM weakness^41^. Abnormal levels of intracellular calcium (Ca^2+^) in the dystrophin-deficient muscle have been reported to contribute to DMD disease progression^42^. To the best of our knowledge, functional DMD-derived NMOs have not been described yet. T-NMOs derived from two DMD patients showed reduced SkM contraction, altered calcium transient and flux upon neuronal stimulation, reproducing key aspects of the disease phenotype. These results further support the idea that our platform could be used to derive patient-specific t-NMO where functional analysis could be done. Dissecting the neural and muscular contribution, investigating early stages of the diseases that precede clinical diagnosis, and testing patient-specific responses to drugs are among some of the exciting opportunities opened by this t-NMO platform.

## Methods

### Human induced pluripotent stem cell derivation and culture

All human iPSCs cell lines in this study were generated by reprogramming in microfluidics^32,43^. For 2D, NMO and t-NMO experiments, hiPSCs were generated from foreskin fibroblast lines (BJ). Three different BJ-derived hiPSC lines were tested. For DMD 1, that carried a stop codon mutation within the exon 61 (c.9100C>T) of the *DMD* gene^33^, primary myogenic cells were purchased from Telethon biobank. For DMD2 and HD, primary skin fibroblasts were derived from a 15-year-old male DMD patient carrying a deletion of exons 45-52 in the *DMD* gene and from a 24-year-old male healthy donor, respectively (ethical committee approval #28062018, University of Milan; #11112021 and #11662022, Policlinico of Milan). Human iPSCs were cultured in feeder free conditions on 0.5% Matrigel (MRF, Corning) coated cell culture plates (6-multiwell, Sarstedt) in StemMACS iPS-Brew XF (Miltenyi Biotec) at 37°C in 5% CO_2_ in cell incubator. All cell lines were tested negative for mycoplasma and maintained below passage 10 before their use for differentiation.

### Decellularized SkMs derivation and characterization of mechanical properties

Murine diaphragms were retrieved from 3 to 5-month-old C57BL/6j mice (protocol N. 1103/2016 and 418/2020-PR approved by Animal wellness local ethics committee, Organismo per il Benessere Animale OPBA, University of Padova and Fondazione Istituto di Ricerca Pediatrica Città della Speranza and Italian Ministry of Health). After collection, diaphragms were washed 2 times in sterile phosphate buffered saline (PBS, Gibco-Fisher Scientific) and then transferred in deionized water in order to start the decellularization process. Diaphragms were processed with 3 detergent-enzymatic treatment (DET) cycles in order to obtain a complete cell removal. Each DET cycle was composed of deionized water at 4 °C for 24 h, 4% sodium deoxycholate (Sigma) at room temperature for 4 h, and 2000 Kunitz DNase-I (Sigma) in 1M NaCl (Sigma) at RT for 3 h^44^. After decellularization, matrices were washed for at least 3 days in PBS and immediately used or preserved in liquid nitrogen.

For atomic force measurements (AFM), freshly isolated murine diaphragm or dSkM were analyzed by using an Atomic Force Microscope (XEBio), mounted on an Inverted Optical Microscope (Nikon Eclipse Ti). This combination enabled positioning of the AFM tip on the area of interest of the samples. All force-displacement curves were collected using PPP-CONTSCR-10 pyramidal tips mounted on Si3N4 cantilevers with a nominal spring constant of 0.2 N/m (NanoSensors). Cantilever spring constants were calibrated by the manufacturer prior to use. Before each test, the sensitivity of the AFM photodetector was calculated by measuring the slope of force-distance curve acquired on a silicon standard. Indentations were performed at a rate of 0.5 μm/s, producing an indentation with a depth of 2 μm. All AFM measurements were done in a fluid environment (PBS) at room temperature. The Young’s modulus was calculated applying a fit of the Hertz model to each individual force curve, assuming a Poisson ratio of 0.5. The moduli of at least 3 samples were calculated as an average over 3-5 sites per sample.

### HiPSC neuromuscular differentiation

Two days before differentiation (Day -2) hiPSCs were enzymatically dissociated as single cells using TryplE Select (Gibco), plated on the specific culture device in StemMACS iPS-Brew XF (Miltenyi Biotec) supplemented with 10 µM StemMACS Y27632 (Miltenyi Biotec), and cultured for 24 h at 37 °C and 5% CO_2_ in cell incubator. One day before differentiation (day -1), media was changed, and cells were cultured in StemMACS iPS-Brew XF. The differentiation protocol started at day 0 and was adapted from literature studies^23^. Briefly, from day 0 to day 2 the media was switched to a Dulbecco’s Modified Eagle Medium/Nutrient Mixture F-12 (DMEM F12, Gibco) based medium, supplemented with Insulin-Transferrin-Selenium (ITS, Gibco), 1% Pen-Strep (Gibco), WNT agonist CHIRON99021 (Miltenyi Biotec) and BMP antagonist LDN193189 (Miltenyi Biotec). From day 3 to day 5, fibroblast growth factor-2 (FGF-2, Immunotools) was supplemented to the media. Starting from day 6, medium was changed to DMEM F-12, supplemented with 15% Knockout™ Serum Replacement (KSR, Gibco), hepatocyte growth factor (HGF, ImmunoTools), insulin-like growth factor 1 (IGF-1, Miltenyi Biotec), FGF-2 and LDN193189. From day 8 to day 11 of differentiation cells were cultured in DMEM F12 supplemented with 15% KSR and IGF-1. From day 12, the previous media was modified by including HGF and IGF-1. From day 26, cells were cultured in myogenic secondary differentiation media composed by DMEM F12 supplemented with KSR, ITS, 1% Pen-Strep and CHIRON99021 until the end point of the experiment (day 30).

For bidimensional differentiation (2D), 0.5% Matrigel-coated cell culture 24-multiwell plates (Sarstedt) were used. Cells were cultured in all the conditions (2D, NMO, t-NMO) following the media changes previously described.

For neuromuscular organoid (NMO) derivation, Matrigel were dispensed on a sterile glass cover slip (Vetrotecnica) forming 1 cm^2^ of Matrigel droplet in 24-multiwell plates (Sarstedt). Droplets were incubated at 37°C for 20 minutes in cell incubator to allow Matrigel polymerization. On day -2, hiPSCs were resuspended in StemMACS iPS-Brew XF supplemented with StemMACS Y27632, seeded on Matrigel droplets and cultured at 37°C and 5% CO_2_ in cell incubator.

For t-NMO derivation, each decellularized murine diaphragm^22^ was surgically dissected into equal parts. On day -2, hiPSCs were resuspended in StemMACS iPS-Brew XF supplemented with StemMACS Y27632, were seeded with dSkMs and cultured at 37°C and 5% CO_2_ in cell incubator.

### Immunofluorescence analysis

T-NMOs were fixed for 1 hour in 4% paraformaldehyde (PFA, Sigma-Aldrich) and NMOs for 1 hour in 2% PFA. Samples were washed twice in phosphate buffered saline (PBS, Sigma-Aldrich) and analyzed in whole mount or in 30 μm cross-sections. For cryosection staining, samples were embedded in optimum cutting temperature (OCT) compound (Sakura 4583) and sectioned with CM1950 cryostat (Leica). Samples were washed with PBS for 5 minutes and blocked at room temperature in 1% bovine serum albumin (BSA, Sigma-Aldrich), 0.5% Triton X-100 (Sigma) in PBS (PBST) solution for 2 hours. Primary antibodies (Supplementary Table 8) were diluted in PBST solution and incubated for 72 hours at 4 °C for whole-mount staining, and overnight at 4 °C for cryosections. Samples were then washed extensively 3 times for 15 minutes each in PBST solution and incubated for 48 hours (whole mount) at 4 °C and 2 hours at room temperature (cryosections) with secondary antibodies solution (Supplementary Table 9). To counterstain the nuclei, 10 µg/mL Hoechst 33342 (ThermoFisher) was used. Samples were washed with PBST before mounting with 80% glycerol (Sigma) in PBS. Images were acquired using LSM800 inverted confocal microscope (Zeiss) and Thunder fluorescent stereomicroscope (Leica M205 FCA) equipped with PLANAP0 1.0X objective.

### Image preparation and analysis

We used ImageJ software for adjustments of levels and contrast, maximum and standard deviation intensity projections, 3D reconstructions and thresholding to create binary masks used for directionality, neuromuscular ratio, myotube and neural projection length and thickness. For quantification, 6 to 10 immunofluorescence images of independent biological triplicates for each sample were converted in binary masks, then analyzed with measurement and directionality plugins of ImageJ software. For distribution of muscular and neural cells in 2D, NMO and t-NMO, spatial maps were obtained by using 3D surface plot plugin of ImageJ software on immunofluorescence images for MYHC/TUJ-1 or desmin/TUJ-1 stained samples. For quantification of NPC and neuron distance from dSkMs, we measured the perpendicular distance to the tangent to the dSkM in cross-sections of t-NMO stained for nuclear markers that identified the cell populations with command distance of ImageJ software. Comparable-sized ROIs were chosen for each of the analyzed images. All mentioned ImageJ plugins have source code available and are licensed under open-source GNU GPL v.3 license.

### Live imaging analysis

To evaluate cell viability in tNMO constructs, samples were treated with calcein (LifeTechnologies). Briefly, samples were washed with PBS and subsequently incubated with a working solution of 3 µM calcein in serum-free cell medium for 30 min at 37°C and 5% CO_2_, and further washed with PBS before imaging. Live imaging analysis was performed using a fluorescent stereomicroscope (Leica M205 FCA) equipped with PLANAP0 1.0X objective.

For calcium flux analysis, 2D and t-NMO samples were treated with Fluo-4-AM (Invitrogen F14201) at day 30 of differentiation. After PBS wash, samples were incubated with 20 µM Fluo-4-AM (Invitrogen), 5 µL/mL Pluronic^TM^ F-127 (Thermo Fisher Scientific), and 12.5 µL/mL sulfinpyrazone (Sigma-Aldrich) in serum-free cell medium for 30 min at 37°C and 5% CO_2_. Samples were then accurately washed with PBS and live imaging was performed with fluorescent stereomicroscope (Leica M205 FCA) equipped with PLANAP0 1.0X objective with an acquisition rate of 16 frame per second. Glu solution was prepared using L-Glutamic acid powder (Sigma-Aldrich) dissolved in sterile water to obtain a 100 mM stock solution. ACh (Sigma-Aldrich) was reconstituted in PBS to produce a 100 mM stock solution. Each neurotransmitter solution was administered during live imaging acquisition at the final working concentration of 10 μM.

For toxin treatments, samples were incubated with 200 pM Botulinum neurotoxin A (BoNT) and 1 µg/ml α-bungarotoxin Alexa Fluor^TM^ 555 (Invitrogen, B13422) for 12 hours at 37°C. BoNT/A was kindly provided by Prof. Cesare Montecucco, University of Padua, IT. After the incubation, samples were analyzed via Fluo-4 live imaging.

### Contraction and calcium analysis

To obtain calcium transient data, fluorescence intensity of chosen regions of interest (ROI) was extracted using the Analyze tool of the Leica Application Suite X (LAS X) software associated with Leica M205 FCA stereomicroscope, according to image acquisition settings. For each category of samples at least 3 independent video acquisition were analyzed, each containing at least 3 ROIs. For every set of data Ca^2+^ dynamics was analyzed using MATLAB 2021a software (MathWorks). For Ca^2+^ peak quantification (peak amplitude), Fluo-4 signals for each ROI were quantified using ImageJ software and reported as relative changes of fluorescence emission intensity (F-F0)/F0, where F0 is the basal fluorescence intensity at time 0 and F is the fluorescence intensity at time t.

To quantify contraction profiles multiple ROI of t-NMO were selected using the Analyze tool of the LAS X software, data were analyzed in MATLAB software.

For each time-lapse, contraction was expressed as power spectrum where the fundamental frequency of the signal was highlighted. A spectrogram was generated to display the contraction pattern of each acquired ROI along the time of acquisition, this allows better visualization and comparison of time-frequency content in the signals.

The power spectrum and the spectrogram were obtained using the Fast Fourier Transform (FFT), an algorithm used to obtain frequency domain data from a signal. For overall contraction of t-NMO before and after toxin treatments, fluorescence intensity variation of entire samples was measured using the Analyze tool of the LAS X software.

### RNA purification and RT-qPCR

Total cell RNA was isolated and purified using RNeasy Plus Mini Kit (Qiagen) according to the manufacturer’s instructions. Briefly, 2D and NMO samples were harvested by direct lysis in the cell culture well. t-NMO samples were accurately disrupted and homogenized with sterile scissors prior to extraction. All the harvested and homogenized samples were lysed with RNeasy Plus lysis buffer (RLT, Qiagen) and further processed according to the protocol. Extracted RNA quality and concentration were assessed with Nanodrop (Thermo Scientific). Complementary DNA (cDNA) of 2D, NMO and t-NMO samples was obtained using High Capacity cDNA Reverse Transcription Kit (Applied Biosystems,) in a dedicated thermocycler (Mastercycler X50a, Eppendorf).

Expression of myogenic and neural markers was quantified by using a 7500 Fast Real-Time PCR System (Applied Biosystem) and Platinum SYBR Green SuperMix kit components (Invitrogen, 11733-038) according to the manufactureŕs instructions. All the primers used are listed in Supplementary Table 10.

Target Ct values of gene expression were normalized to that of myosin heavy chain, used as an housekeeping gene. Data are shown as relative fold change to 2D gene expression applying the 2^−ΒΒCT^ method.

### Bulk sequencing analysis

Total RNA was quantified using the Qubit 4.0 fluorometric Assay (Thermo Fisher Scientific). Libraries were prepared from 125 ng of total RNA using the NEGEDIA Digital mRNA-seq research grade sequencing service (Next Generation Diagnostic srl)^46^ which included library preparation, quality assessment and sequencing on a NovaSeq 6000 sequencing system using a single-end, 100 cycle strategy (Illumina Inc.).

The raw data were analyzed by Next Generation Diagnostic srl proprietary NEGEDIA Digital mRNA-seq pipeline (v2.0) which involves a cleaning step by quality filtering and trimming, alignment to the reference genome and counting by gene^47,48^.

Bioinformatic analysis was performed in R v. 4.1.1 with Bioconductor v. 3.^49^. Genes were annotated using R package org.Hs.eg.db v. 3.13. Genes were filtered out if not having at least 2 replicates from the same culture condition with at least 2 raw counts. Raw counts were subsequently normalized and expressed as counts per million (CPM) using R package DESeq2 v. 1.^50^. Log2 scale CPM data were obtained adding a unit pseudo-count to avoid infinite values. PCA was performed by R stats package function *prcomp* by singular value decomposition (SVD) after centering. Differentially expressed genes (DEGs) were computed using DESeq2 starting from raw count data, using a *P* value, after correction by the Benjamini Hochberg method, lower than 0.01. DEGs between t-NMO samples and 2D or NMO conditions were input for an overrepresentation analysis within Gene Ontology (GO, https://www.ebi.ac.uk/GOA) and Reactome (https://reactome.org/) databases. The analysis was performed using ClueGO v. 2. 5.^51^ within Cytoscape v. 3.8.^52^ environment for network visualization. A Benjamini-Hochberg corrected *P* value cutoff of 0.0001 was set. Hierarchical clustering with heat map visualization was performed using R package pheatmap v. 1.0.12, using Euclidean distance.

### Statistical analysis

All analyses were performed with GraphPad prism 6. We expressed data as mean±s.e.m or mean±s.d of multiple biological replicates (as indicated in the figure legend). We determined statistical significance by unequal variance Student’s t-test, one-way analysis of variance (ANOVA) and Tukey’s multiple comparison test are shown in Supplementary Tables 1,4,5,7. A P value of less than 0.05 was considered statistically significant.

## Acknowledgments

This work was supported by STARS Starting Grant 2017 of University of Padova (grant code LS3-19613), by Bando Direzione Scientifica IRP Città della Speranza (grant code 21/05) by BIRD (code URCI_BIRD2223_01) to A.U.; by AFM Telethon (grant code) to N.E.; and by Bando CARIPARO-Ricerca Pediatrica 2020-2022 (grant number 20/17 FCR) to P.M. O.G. was supported by University of Padova under the 2019 STARS Grants program (iNeurons). DMD1 cell lines were provided by the Bank of muscle tissue, peripheral nerve, DNA and Cell Culture, member of Telethon network of Genetic biobanks, at Fondazione IRCCS Ca’ Granda, Ospedale Maggiore Policlinico, Milano, Italy”. We thank Marco Braggion and Marco La Placa for helping with MATLAB software.

## Author contributions

A.U. designed the study. B.A. performed the differentiation experiments and analysis. F.C, L.R., G.B., P.C., A.L., contributed to sample preparation and imaging analysis. A.U., B.A., L.R, F.C., G.C., and A.L. performed the live imaging analysis. P.R. and M.E. helped in setting the seeding strategies. O.G., S.A. and W.Q performed cell reprogramming. C.La. contributed to neural characterization and data interpretation. E.M. and M.P. derived dSkM. M.G. performed AFM analysis and data interpretation. D.C. performed RNA-sequencing. M.C. and S.C. contributed to NMJ imaging analysis. C.Lu. analyzed RNA-seq data. C.V. and Y.T provided the DMD2 and HD primary cells. N.E. supervised the reprogramming and mechanical properties investigation, and critically contributed to the transcriptomic analysis and to data interpretation. All the authors contributed to revision of the manuscript. A.U. analyzed and interpreted the data, wrote the manuscript and supervised the project.

