## Supplementary Material for "Native extracellular matrix promotes human neuromuscular organoid morphogenesis and function"

**Supplementary Figures:**

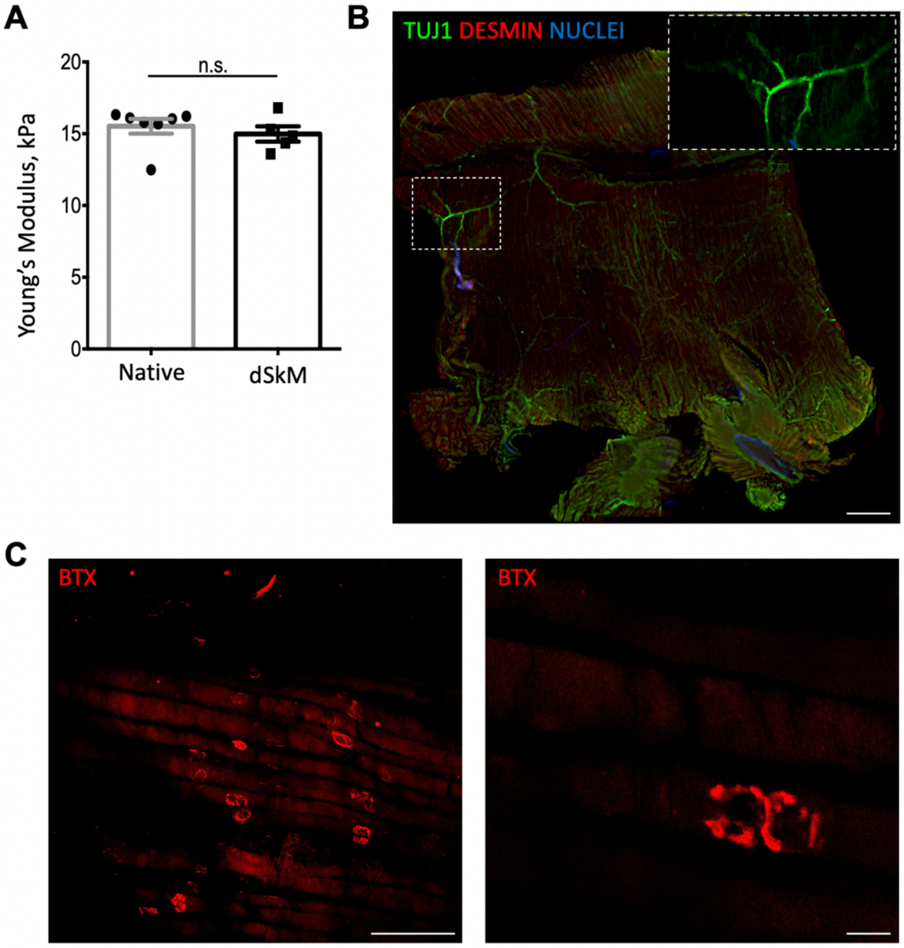

**Supplementary Figure 1.**
**Characterization of dSkM. A.** Young's modulus of native freshly isolated murine diaphragm (native, grey) and decellularized murine diaphragm (dSkM, black) measured by atomic force microscopy. Data are shown as mean ± s.d. of 5 or 7 independent replicates; unequal variance Student's t-test; n.s., not statistically significant. **B.** Representative image showing whole mount immunofluorescence staining for TUJ-1 (green), Desmin (red), Nuclei (blue) of dSkM. The inset shows a higher magnification of a branched TUJ-1 structure. Scale bar, 500 μm. **C.** Representative fluorescent images showing lower (left) and higher (right) magnification of dSkM whole mount stained with bungarotoxin (BTX, red). Scale bars, 100 μm (left) and 10 μm (right).

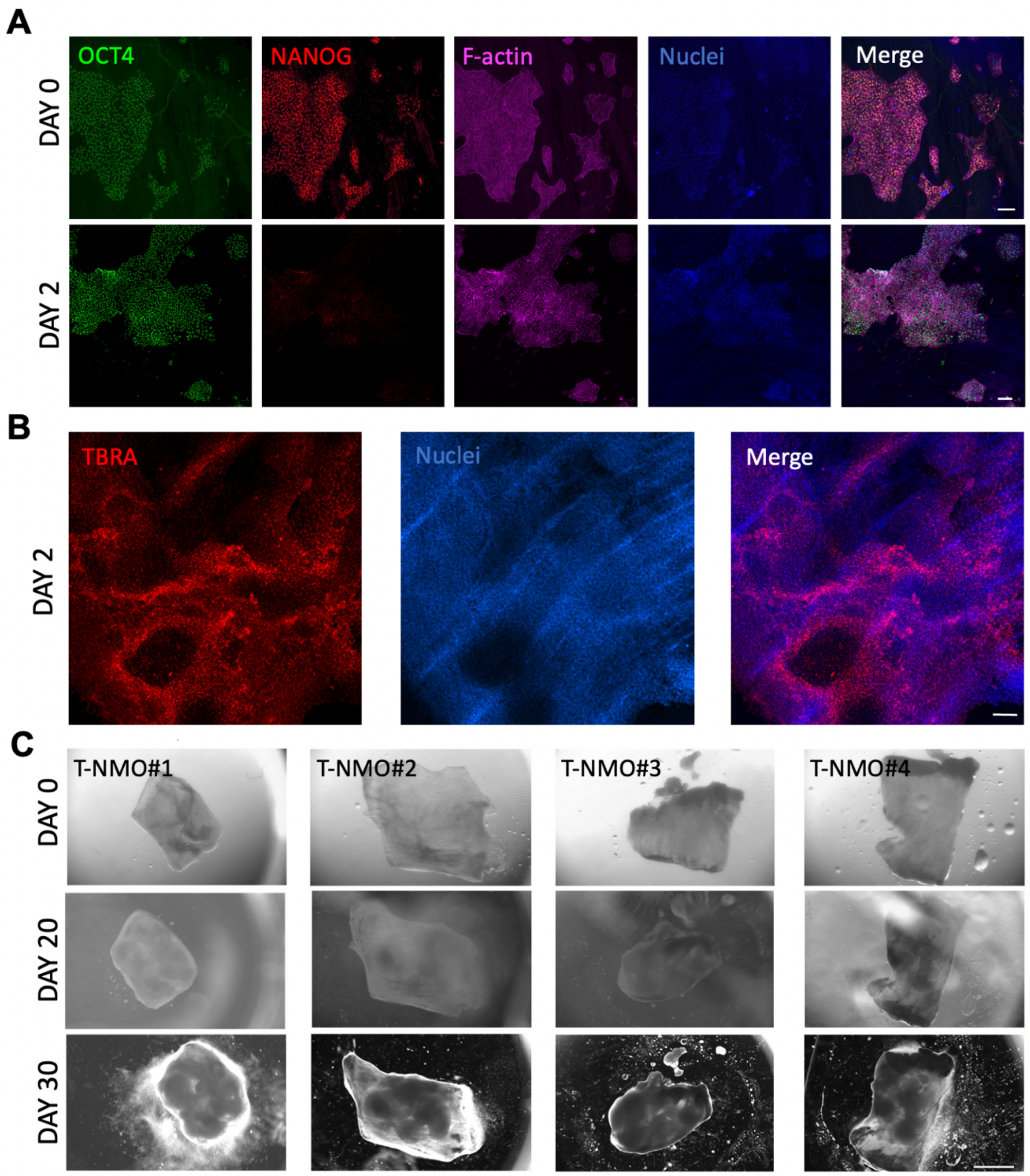

**Supplementary Figure 2.**
**Characterization and reproducibility of t-NMO production. A.** Representative images showing immunofluorescence staining for OCT4 (green), NANOG (red), F-actin (far-red), Nuclei (blue) of t-NMO at early time points. The insets show higher magnifications. Scale bars, 100 µm. **B.** Representative images showing immunofluorescence staining for Brachyury (TBRA, red), Nuclei (blue) of t-NMO at day 2 of hiPSC differentiation. Scale bars, 100 µm. **C.** Bright field images showing development of cell populations in four different t-NMO samples. Scale bars, 2 mm.

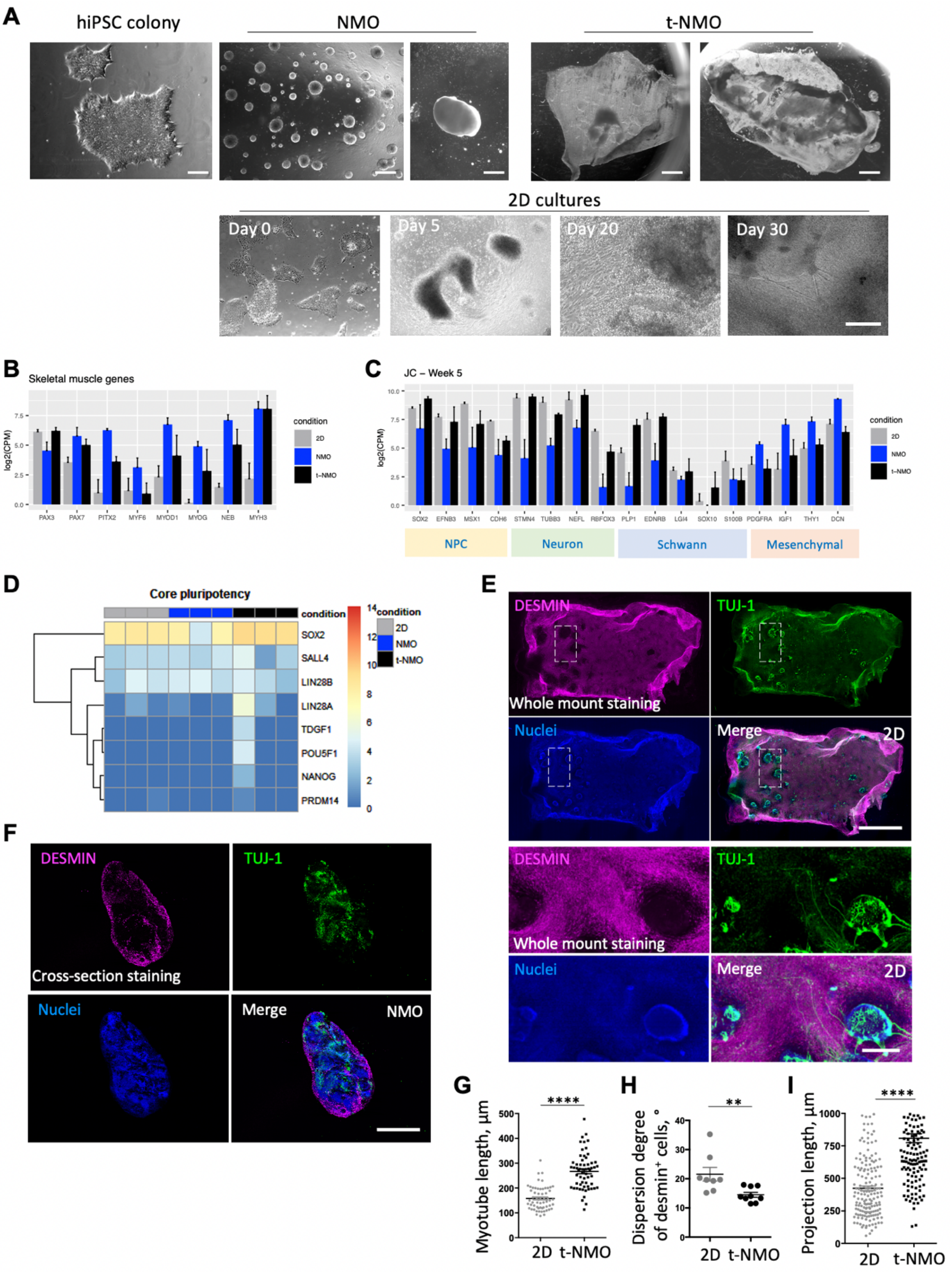

**Supplementary Figure 3.**

**Neuromuscular differentiation can be achieved in 2D cultures, and in 3D producing NMO and t-NMO after 30 days of hiPSC differentiation. A.** Representative bright field images showing morphologic appearance of hiPSCs colonies before (left upper panel) or 5 and 30 days after differentiation onto Matrigel (NMO, middle upper panels) or dSkM (t-NMO, right upper panels). Lower panels show hiPSCs seeded (day 0) into conventional cell culture plates (2D) undergoing toward neuromuscular differentiation after 5 (Day 5), 20 (Day 20) or 30 days (Day 30) from the treatment. Scale bars, 20 µm hiPSC colony panel, 100 µm (left) and 1 mm (right) for NMO panels, 2 mm for t-

NMO panels, 50 μm for 2D panels. **B.** Expression profile from bulk RNA-seq measurement of SkM genes. **C.** Expression profile from bulk RNA-seq measurement of genes typically of neural progenitor cells (NPC), neurons, Schwann and mesenchymal cells. **D.** Hierarchical clustering with heat map visualization of core pluripotency genes. Color bar represents non-centred log2(CPM). **E.** Representative immunofluorescent images showing myogenic (desmin, magenta) and neuronal (TUJ-1, green) cells in whole mount stained 2D cultures. The inset for 2D cultures shows a higher magnification field represented in the lower panels. Scale bars, 1 cm (upper), 20 mm (lower). **F.** Representative immunofluorescent images showing myogenic (desmin, magenta) and neuronal (TUJ-1, green) cells in in cryo-sectioned NMO (lower panels). Scale bar, 1 mm. **G.** Quantification of desmin$^{+}$ myotube length derived from hiPSCs after 30 days of differentiation in 2D cultures (gray) or t-NMO (black). Data are shown as mean ± SEM of 3 independent replicates; dots represent single measured myotubes; unequal variance Student's *t*-test was used. ****$P < 0.001$. **H.** Quantification of dispersion degree of desmin$^{+}$ myotube directionality derived from hiPSCs after 30 days of differentiation in 2D cultures (gray) or t-NMO (black). Data are shown as mean ± SEM of 3 independent replicates; dots represent single measured ROIs; unequal variance Student's *t*-test was used. **$P = 0.0095$. **I.** Quantification of TUJ-1$^{+}$ neural projection length derived from hiPSCs after 30 days of differentiation in 2D cultures (gray) or t-NMO (black). Data are shown as mean ± SEM of 3 independent replicates and dots represent single measured neural projections; unequal variance Student's *t*-test was used. ****$P < 0.001$.

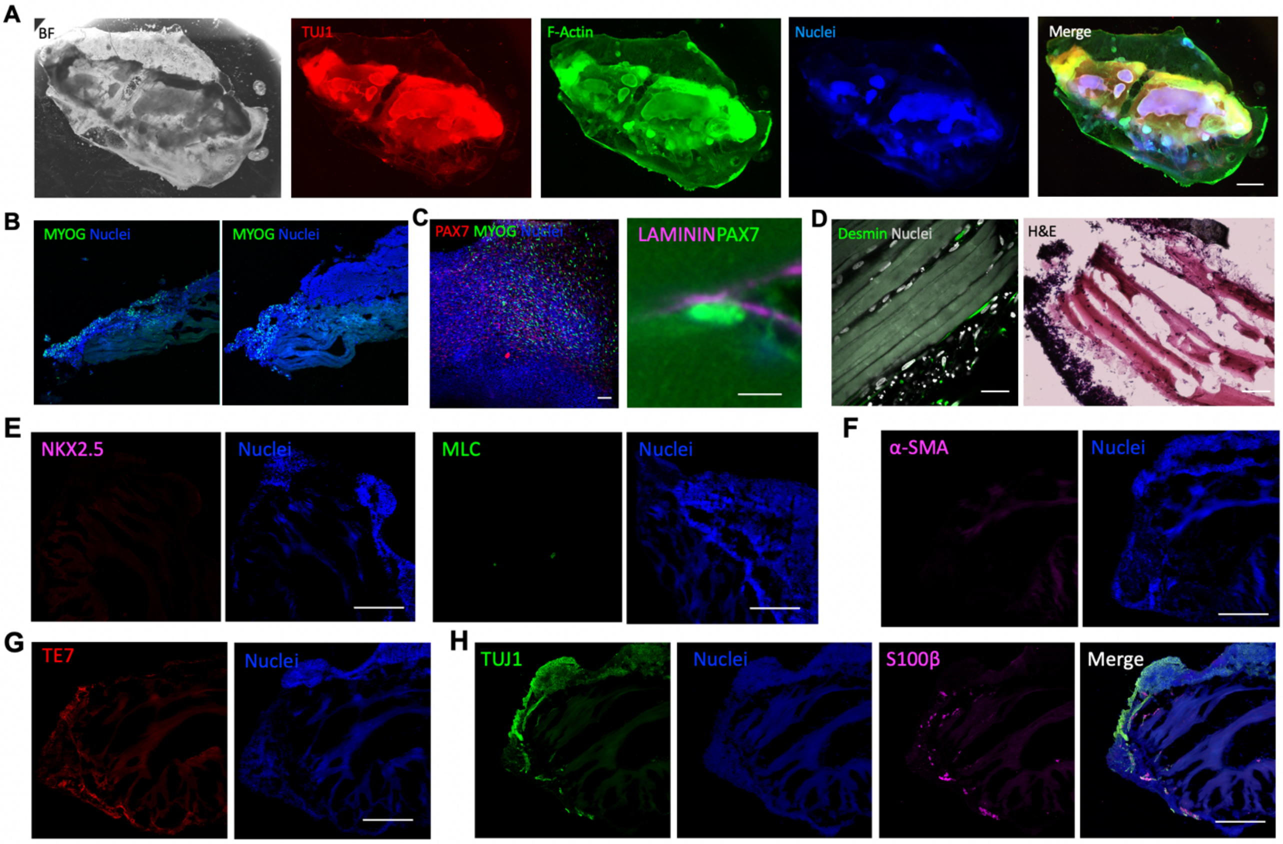

**Supplementary Figure 4.**

**Immunofluorescence characterization of t-NMO after 30 days of differentiation. A.** Representative stereomicroscope images showing bright field (BF) and immunofluorescence staining for TUJ-1 (red), F-actin (green) and nuclei (blue) of t-NMO at day 30 of differentiation. Nuclei were stained with Hoechst (blue). Scale bar, 1 mm**. B.** Representative images showing immunofluorescence staining for MYOG (green) of t-NMO cryo-sections. Nuclei were stained with Hoechst (blue). Scale bar, 200 μm. **C.** Left, representative wholemount imaging showing PAX7 and MYOG expressing cells in t-NMOs. Right, representative confocal Z-stack image showing immunofluorescence staining for PAX7 (green) and laminin (magenta). Scale bar, 10 μm. **D.** Representative images showing cell invasion within the dSkM of t-NMO at day 30 of differentiation. Left, whole mount immunofluorescence staining showed myogenic cells expressing Desmin (green) located on the top and within the inner area of dSkMs. Nuclei were stained with Hoechst (blue). Scale bars, 50 μm. Right, hematoxylin-eosin (H&E) staining of t-NMO cryosections. Scale bar, 100 μm. **E.** Representative images of t-NMO cross-section showing immunofluorescence staining for cardiac markers homeobox protein NKX2.5 (magenta, left) and myosin light chain (MLC, green, right). Nuclei were stained with Hoechst (blue). Scale bar, 200 μm. **F.** Representative images of t-NMO cross-section showing immunofluorescence staining for alpha-smooth muscle actin (α-SMA). Nuclei were stained with Hoechst (blue). Scale bar, 200 μm. **G.** Representative images of t-NMO cross-section showing immunofluorescence staining for fibroblast marker TE7 (red). Nuclei were stained with Hoechst (blue). Scale bar, 200 μm. **H.** Representative images showing immunofluorescence staining for TUJ-1 (green), S100 calcium-binding protein B (S100β, magenta) Schwann cells marker. Nuclei were stained with Hoechst (blue). Scale bar, 200 μm.

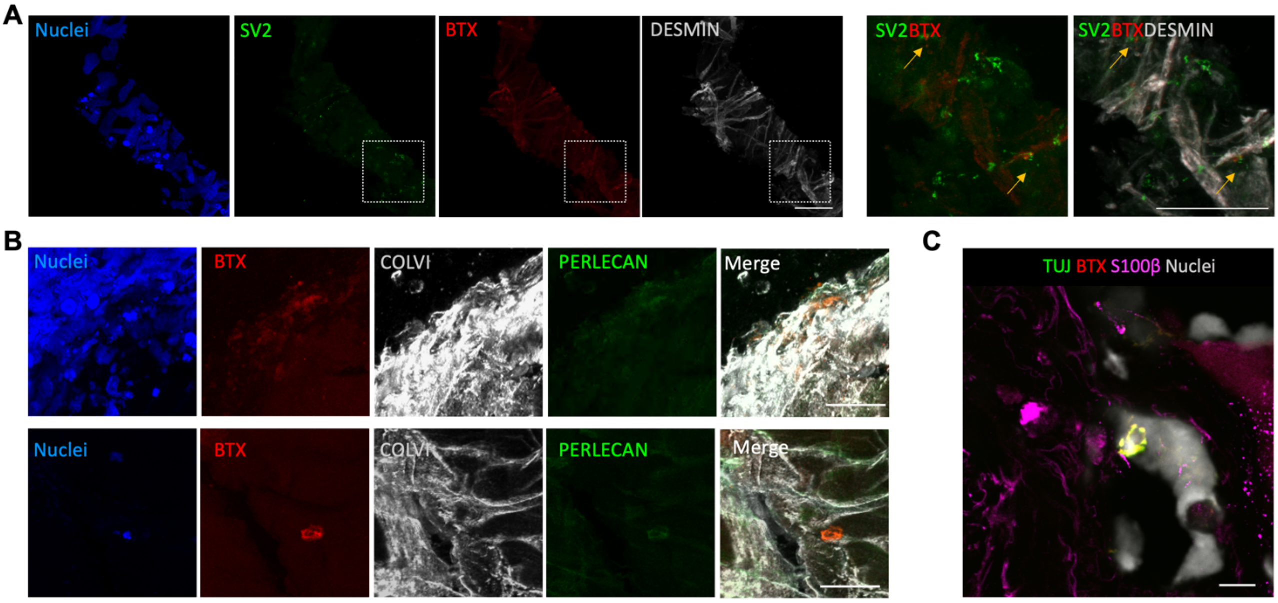

**Supplementary Figure 5.**

**NMJ characterization in t-NMOs. A.** Representative immunofluorescence images showing SV2 (green) and BTX (red) NMJ compartments, and desmin (grey) organization (right panel) and the inset (left panel) showing higher magnification and juxtaposition of the SV2 and BTX regions (yellow arrows). Nuclei were counterstained with Hoechst (blue). Scale bars, 12.5 µm (left) and 25 µm (right). **B.** Representative immunofluorescence images showing BTX (red) regions localization within ECM stained for collagen VI (COLVI, grey) and proteoglycans (PERLECAN, green). Nuclei were stained with Hoechst (blue). Scale bar, 25 µm. **C.** Representative image of immunofluorescence staining for TUJ-1 (green), BTX (red) and S100β (far-red). Nuclei were stained with Hoechst (grey). Scale bar, 10 µm.

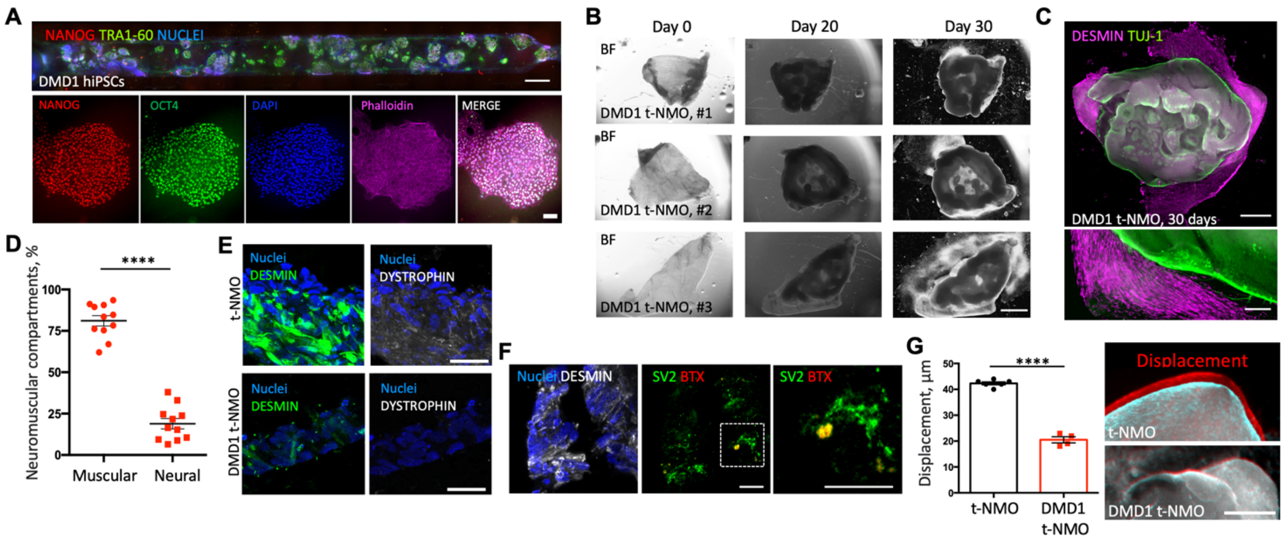

**Supplementary Figure 6.**

**DMD1 t-NMO derivation and characterization. A.** Representative images showing immunofluorescence staining for NANOG (red), TRA1-60 (green) and Nuclei (blue) (upper panel, scale bar, 1 mm) and OCT4 (green), Nanog (red), F-actin (magenta), Nuclei (blue) (lower panel, scale bar, 100 µm) of DMD1 hiPSCs. **B.** Bright field images showing morphological appearance of four different t-NMO samples at time 0 (Day 0) of differentiation and after 20 or 30 days of differentiation. Scale bar, 2 mm **C.** Representative stereomicroscope images of whole mount immunofluorescence staining for TUJ-1 (green) and desmin (magenta) of DMD1 t-NMO after 30 days of differentiation. Scale bar, 1 mm (upper panel). The lower panel shows higher magnification, scale bar 200 µm. **D.** Quantification of the area occupied by muscular (Desmin$^{+}$) or neuronal (TUJ-1$^{+}$) cells derived from DMD1 hiPSCs after 30 days of differentiation into DMD1 t-NMO. Data are shown as mean ± SEM of 10 ROIs of 3 independent replicates; unequal variance Student's *t*-test was used. ****$P < 0.001$. **E.** Representative immunofluorescence images showing desmin (green) and Dystrophin (grey) in cryosection of t-NMO (upper panels) and DMD1 t-NMO (lower panels). Nuclei were stained with Hoechst (blue). Scale bar, 25 µm. **F**. Representative immunofluorescence images showing SV2 (green) and BTX (red) pre- and post-NMJ compartments (middle panel), in desmin (grey) expressing area (left panel). The inset (right) shows higher magnification and juxtaposition of the SV2 and BTX regions (yellow arrows). Nuclei were counterstained with Hoechst (blue). Scale bars, 25 µm. **G.** Quantification of t-NMO and DMD1 t-NMO displacement expressed in µm, upon Glu stimulation (left panel). Representative stereomicroscope images showing the displacement (red) of t-NMO (upper panel) and DMD1 t-NMO (lower panel) upon Glu stimulation. Data are shown as mean ± SEM of 4 independent replicates; unequal variance Student's *t*-test was used. ****$P < 0.001$. Scale bar 500 µm.

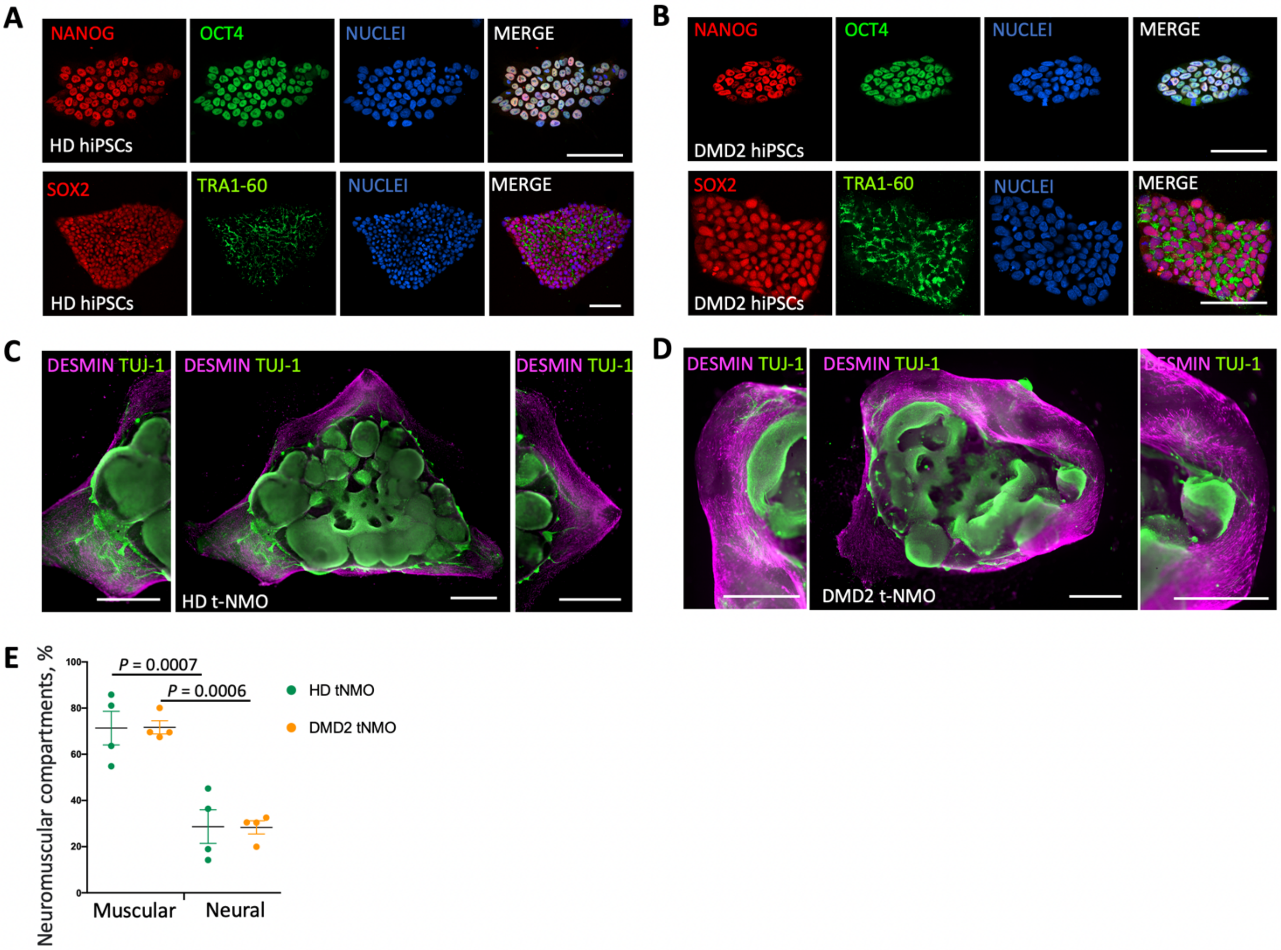

**Supplementary Figure 7.**

**DMD2 and HD t-NMO derivation and characterization. A.** Representative images showing immunofluorescence staining of HD hiPSCs for NANOG (red), OCT4 (green) and Nuclei (blue) (upper panels) and SOX2 (red), TRA1-60 (green), Nuclei (blue) (lower panels). Scale bars, 100 μm. **B.** Representative images showing immunofluorescence staining of DMD2 hiPSCs for NANOG (red), OCT4 (green) and Nuclei (blue) (upper panels) and SOX2 (red), TRA1-60 (green), Nuclei (blue) (lower panels). Scale bars, 100 μm. **C.** Representative stereomicroscope images of whole mount immunofluorescence staining for TUJ-1 (green) and desmin (magenta) of HD t-NMO at day 30 of differentiation (middle). The lateral panels show higher magnification. Scale bars, 1 mm. **D.** Representative stereomicroscope images of whole mount immunofluorescence staining for TUJ-1 (green) and desmin (magenta) of DMD2 t-NMO at day 30 of differentiation (middle). The lateral panels show higher magnification. Scale bars, 1 mm. **E.** Quantification of the area occupied by muscular (Desmin$^{+}$) or neuronal (TUJ-1$^{+}$) cells derived from HD or DMD2 hiPSCs after 30 days of differentiation into HD (green) or DMD2 t-NMO (orange), respectively. Data are shown as mean ± SEM of 4 ROIs; One-way ANOVA with Tukey's multiple comparisons test.

**Supplementary Table 1. Statistical analysis of Figure 1E.**

| Tukey's multiple comparisons test | Significant | Summary | Adjusted P Value |
|---|---|---|---|
| **Figure 1E** | | | |
| 2D muscular vs. NMO muscular | No | ns | 0.7804 |
| 2D muscular vs. t-NMO muscular | No | ns | 0.1142 |
| 2D muscular vs. 2D neural | Yes | **** | <0.0001 |
| 2D muscular vs. NMO neural | Yes | **** | <0.0001 |
| 2D muscular vs. t-NMO neural | Yes | **** | <0.0001 |
| NMO muscular vs. t-NMO muscular | No | ns | 0.9285 |
| NMO muscular vs. 2D neural | Yes | **** | <0.0001 |
| NMO muscular vs. NMO neural | Yes | **** | <0.0001 |
| NMO muscular vs. t-NMO neural | Yes | **** | <0.0001 |
| t-NMO muscular vs. 2D neural | Yes | **** | <0.0001 |
| t-NMO muscular vs. NMO neural | Yes | **** | <0.0001 |
| t-NMO muscular vs. t-NMO neural | Yes | **** | <0.0001 |
| 2D neural vs. NMO neural | No | ns | 0.7804 |
| 2D neural vs. t-NMO neural | No | ns | 0.1142 |
| NMO neural vs. t-NMO neural | No | ns | 0.9285 |
| **Figure 1J** | | | |
| 2D vs. NMO | Yes | **** | <0,0001 |
| 2D vs. t-NMO | Yes | **** | <0,0001 |
| NMO vs. tNMO | Yes | **** | <0,0001 |
| **Figure 1K** | | | |
| 2D vs. NMO | Yes | **** | <0,0001 |
| 2D vs. tNMO | Yes | **** | <0,0001 |
| NMO vs. tNMO | Yes | **** | <0,0001 |

**Supplementary Table 2.**

**Results of bulk RNA-seq differential expression analysis of 2D, NMO, and t-NMO samples.** Each sheet reports the list of DEGs obtained from the comparison shown in the tab name, and the normalized CPM in each sample.

**Supplementary Table 3.**

**Results of bulk RNA-seq enrichment analysis.** Each sheet reports the enriched categories from Reactome and GO databases, which were obtained in the analysis of DEGs between t-NMO samples compared to 2D and NMO conditions, respectively.

**Supplementary Table 4. Statistical analysis of Figure 4C.**

| | Significant | Summary | Adjusted P Value |
|---|---|---|---|
| 2D vs. NMO | Yes | *** | 0.0008 |
| 2D vs. t-NMO | Yes | **** | <0.0001 |
| NMO vs. t-NMO | Yes | **** | <0.0001 |

**Supplementary Table 5. Statistical analysis of Figure 5D.**

| **Tukey's multiple comparisons test** | **Significant** | **Summary** | **Adjusted P Value** |
|---|---|---|---|
| CTRL 0H vs. CTRL 24H | No | ns | 0,947 |
| CTRL 0H vs. BONT 0H | No | ns | 0,3456 |
| CTRL 0H vs. BONT 12H | Yes | * | 0,0176 |
| CTRL 0H vs. BTX 0H | No | ns | 0,7442 |
| CTRL 0H vs. BTX 12H | Yes | ** | 0,0079 |
| CTRL 24H vs. BONT 0H | No | ns | 0,0943 |
| CTRL 24H vs. BONT 12H | No | ns | 0,0758 |
| CTRL 24H vs. BTX 0H | No | ns | 0,2897 |
| CTRL 24H vs. BTX 12H | Yes | * | 0,0339 |
| BONT 0H vs. BONT 12H | Yes | *** | 0,0006 |
| BONT 0H vs. BTX 0H | No | ns | 0,9729 |
| BONT 0H vs. BTX 12H | Yes | *** | 0,0003 |
| BONT 12H vs. BTX 0H | Yes | ** | 0,0019 |
| BONT 12H vs. BTX 12H | No | ns | 0,996 |
| BTX 0H vs. BTX 12H | Yes | *** | 0,0009 |

**Supplementary Table 6.**
**Results of bulk RNA-seq differential expression analysis of t-NMO vs. t-NMO DMD samples.** Each sheet reports the list of DEGs up- or down- regulated, as indicated in the tab name, and the normalized CPM in each sample.

**Supplementary Table 7. Statistical analysis of Figure 5I.**

| **Glu Stimulation** | | | |
|---|---|---|---|
| **Tukey's multiple comparisons test** | **Significant** | **Summary** | **Adjusted P Value** |
| t-NMO vs. HD t-NMO | No | ns | 0,6534 |
| t-NMO vs. DMD1 t-NMO | Yes | * | 0,041 |
| t-NMO vs. DMD2 t-NMO | Yes | ** | 0,0038 |
| HD t-NMO vs. DMD1 t-NMO | Yes | * | 0,0167 |
| HD t-NMO vs. DMD2 t-NMO | Yes | ** | 0,0023 |
| DMD1 t-NMO vs. DMD2 t-NMO | No | ns | 0,6624 |
| **ACh Stimulation** | | | |
| **Tukey's multiple comparisons test** | **Significant** | **Summary** | **Adjusted P Value** |
| t-NMO vs. HD t-NMO | No | ns | 0,169 |
| t-NMO vs. DMD1 t-NMO | Yes | *** | 0,0002 |
| t-NMO vs. DMD2 t-NMO | Yes | ** | 0,005 |
| HD t-NMO vs. DMD1 t-NMO | Yes | **** | < 0,0001 |
| HD t-NMO vs. DMD2 t-NMO | Yes | *** | 0,0007 |
| DMD1 t-NMO vs. DMD2 t-NMO | No | ns | 0,3842 |

**Supplementary Table 8.** List of primary antibodies used in this study.

| Antibody | Host | Dilution | Company |
|---|---|---|---|
| Adult Myosin Heavy Chain (MYHC) | Mouse | 1:25 | DSHB |
| Alfa-Smooth Muscle Actin (α-SMA) | Mouse | 1:100 | Abcam |
| BAD5 Myosin Heavy Chain 1 (SlowMHC) | Mouse | 1:50 | DSHB |
| Brachyury (TBRA) | Goat | 1:40 | Biotechne |
| Caudal type homeobox 2 (CDX2) | Mouse | 1:100 | Santa Cruz Biotechnology |
| Collagen VI (COLVI) | Rabbit | 1:1000 | Fitgerald Ind. |
| DESMIN | Rabbit | 1:300 | Abcam |
| DESMIN | Mouse | 1:75 | Agilent DAKO |
| DYSTROPHIN | Rabbit | 1:50 | Santa Cruz Biotechnology |
| Homeobox transcription factor (HB9) | Mouse | 1:25 | DSHB |
| Homeobox transcription factor NANOG | Rabbit | 1:100 | Cell signaling |
| ISLET1/2 | Mouse | 1:25 | DSHB |
| Ki67 | Rabbit | 1:200 | Abcam |
| Laminin (LAM) | Rabbit | 1:300 | Sigma-Aldrich |
| Microtubule associated protein 2 (MAP2) | Chicken | 1:10000 | Abcam |
| Myogenin (MYOG) | Rabbit | 1:25 | Santa Cruz Biotechnology |
| Myosin Light Chain 2 (MLC) | Rabbit | 1:200 | Abcam |
| Neuron-specific class III beta-tubulin (TUJ1) | Mouse | 1:5000 | Biolegend |
| Neuronal nuclear antigen (NEUN) | Rabbit | 1:500 | Abcam |
| NK2 Homeobox 5 (NKX2.5) | Goat | 1:200 | Abcam |
| Octamer - binding transcription factor 4 (OCT4) | Mouse | 1:200 | Santa Cruz Biotechnology |
| Oligodendrocyte transcription factor (Olig2) | Mouse | 1:25 | DSHB |
| Paired box protein (PAX7) | Mouse | 1:25 | DSHB |
| PERLECAN | Rat | 1:50 | Santa Cruz Biotechnology |
| Podocalyxin (TRA1-60) | Mouse | 1:100 | Cell signaling |
| S-100 protein beta chain (S100β) | Rabbit | 1:100 | Abcam |
| Sc-71 Myosin Heavy Chain 2 (FastMHC) | Mouse | 1:50 | DSHB |
| SRY – Box transcription factor 2 (SOX2) | Goat | 1:50 | Biotechne |
| SRY – Box transcription factor 2 (SOX2) | Rabbit | 1:500 | Millipore |
| Synaptic vesicle glycoprotein 2 (SV2) | Mouse | 1:50 | DSHB |
| TE7 | Mouse | 1:100 | Millipore |

**Supplementary Table 9.** List of secondary antibodies and fluorescent dyes used in this study.

| Antibody | Host | Dilution | Company |
|---|---|---|---|
| anti-rabbit 488 | Donkey | 1:500 | Invitrogen |
| anti-rabbit 594 | Donkey | 1:500 | Invitrogen |
| anti-rabbit 647 | Donkey | 1:500 | Life Tech |
| anti-mouse 488 | Donkey | 1:500 | Invitrogen |
| anti-mouse 594 | Donkey | 1:500 | Invitrogen |
| anti-mouse 647 | Donkey | 1:500 | Life Tech |
| anti-goat 488 | Donkey | 1:500 | Life Technologies |
| anti-chiken | Chicken | 1:500 | Abcam |
| Phalloidin (F-Actin) 488 | | 1:300 | Invitrogen |
| Phalloidin (F-Actin) 647 | | 1:300 | Invitrogen |
| Bungarotoxin 555 | | 1:1000 | Life Technologies |
| Bungarotoxin 647 | | 1:1000 | Life Technologies |

**Supplementary Table 10. List of primers used in this study.** Oligonucleotide sequences specific for the different genes of interest were used to quantify gene expression of epsilon and gamma AchR expression. Myosin heavy chain was used as housekeeping gene.

| Gene | Primer probe | Sequence | References |
|---|---|---|---|
| Acetylcholine Receptor Epsilon (ACHRE) | Forward (5'->3')<br>Reverse (3'->5') | GCCTGAGGATACTGTCACCATC<br>GTCCTTGCTGTAGTTGAGTCGG | OriGene NM_000080 |
| Acetylcholine Receptor Gamma (ACHRG) | Forward (5'->3')<br>Reverse (3'->5') | CTGTCTTCCTCTTCCTTGTGGC<br>CGACAATGAGGATGGTCACCAC | OriGene NM_005199 |
| Myosin Heavy Chain (MyHC,pan) | Forward (5'->3')<br>Reverse (3'->5') | CAGCCTGGAGCAGCTGTGCAT<br>ATGCCCATAGGCTTCTCGATGAGCTC | Ref. n. 15 |

**Supplementary Video 1. Live imaging of spontaneous contraction relative to Figure 4A.** Representative time-lapse stereomicroscope analysis of alive t-NMO cells (calcein stained, green) that reveal spontaneous rhythmic contraction of t-NMO after 15 days of differentiation.

**Supplementary Video 2. Live imaging showing calcium transients before and after Glutamate and Acetylcholine administration relative to Figure 5.** Representative Fluo-4 calcium dye time-lapse stereomicroscope analysis of t-NMO after 30 days of differentiation. Spontaneous or induced stimulation of neural (Glutamate) or muscular (Acetylcholine) compartments are indicated during the movie.
